# Interferon-armed RBD dimer enhances the immunogenicity of RBD for sterilizing immunity against SARS-CoV-2

**DOI:** 10.1101/2021.05.12.443228

**Authors:** Shiyu Sun, Yueqi Cai, Tian-Zhang Song, Yang Pu, Lin Cheng, Hairong Xu, Chaoyang Meng, Yifan Lin, Jing Sun, Silin Zhang, Yu Gao, Jian-Bao Han, Xiao-Li Feng, Dan-Dan Yu, Yalan Zhu, Pu Gao, Haidong Tang, Jincun Zhao, Zheng Zhang, Jiaming Yang, Zenxiang Hu, Yang-Xin Fu, Yong-Tang Zheng, Hua Peng

## Abstract

Severe acute respiratory syndrome coronavirus 2 (SARS-CoV-2) has caused a global crisis, urgently necessitating the development of safe, efficacious, convenient-to-store, and low-cost vaccine options. A major challenge is that the receptor-binding domain (RBD)-only vaccine fails to trigger long-lasting protective immunity if used solely for vaccination. To enhance antigen processing and cross-presentation in draining lymph nodes (DLNs), we developed an interferon (IFN)-armed RBD dimerized by immunoglobulin fragment (I-R-F). I-R-F efficiently directs immunity against RBD to DLN. A low dose of I-R-F induces not only high titer long-lasting neutralizing antibodies but also comprehensive T cell responses than RBD, and even provides comprehensive protection in one dose without adjuvant. This study shows that the I-R-F vaccine provides rapid and complete protection throughout upper and lower respiratory tracts against high dose SARS-CoV-2 challenge in rhesus macaques. Due to its potency and safety, this engineered vaccine may become one of the next-generation vaccine candidates in the global race to defeat COVID-19.

## Introduction

The pandemic of COVID-19 caused by Severe Acute Respiratory Syndrome Coronavirus 2 (SARS-CoV-2) has swept across the world since its outbreak in late 2019^1^. Mutant coronaviruses continue to evolve, some with improved receptor-binding affinity and infectivity. Although various categories of vaccine candidates against SARS-CoV-2 have been developed, improved vaccines are still urgently needed for public health and various socio-economic crises. In addition to its long-lasting potency, vaccines should be stable in 4-25°C storage, easy to produce, low-cost, and safe for all ages to be practical for the hardest hit, densely populated, and under-resourced countries around the world.

The leading vaccine candidates are mRNA-based, inactivated, or adenovirus-based vaccines (https://vac-lshtm.shinyapps.io/ncov_vaccine_landscape/). Inactivated vaccines can be made by traditional methods. Antibodies induced by such inactivated vaccines target all viral proteins, mostly not related to neutralization. Natural monomeric S protein or RBD yields low titers of neutralizing antibodies due to the poor immunogenicity ^2-4^. Some studies showed that modified S or RBD such as S-trimer and RBD-dimer had been developed to generate higher neutralizing antibodies than monomeric proteins^5, 6^. Also, RBD fused with the Fc domain showed potential immune effects better than RBD^7-9^. Alum adjuvant is commonly used in these viral-antigen vaccines to induce stronger humoral immunity preferentially, but not to be favorable to the T cell responses, especially the type 1 T helper (Th1) cell and cytotoxic T lymphocyte responses (CTLs)^2-4^. Although novel adjuvants may be more potent, they are challenging to prepare and increase the risk of severe side effects with limited usage^5, 10, 11^. Recombinant vaccines based on adenovirus (AdV) vectors, such as Ad5-nCoV, stimulated both B-cell and T-cell responses. However, the ubiquitously pre-existing anti-vector immunity may wreck immune responses resulting in low neutralization antibody titers in trials and invalid immune boost after repeated vaccination^12-14^. The mRNA-based vaccines are the current leading vaccines due to rapid manufacturability after new outbreaks and induce moderate to strong antibody responses and T cell responses. However, it remains to be determined if the reactogenicity of certain mRNA vaccines varies by age and race. Strict conditions for preservation and transportation of mRNA vaccines further limit its broader applicability, especially in developing countries^15-18^.

To increase the immunogenicity of RBD, we have developed a next-generation fusion-protein vaccine, named I-R-F: RBD is armed with an interferon-α (IFNα) at the N-terminus and dimerized by human IgG1 Fc at the C-terminus to target and alert dendritic cells in lymph nodes (LNs). Armed with IFN and dimerized by Fc, which enhances antigen processing and presentation, low dose I-R-F showed more potent immunogenicity than monomeric RBD, inducing robust antibody titers of balanced IgG1 and IgG2a subtypes and robust CD8^+^ T cell response, even without additional adjuvant. We further added a pan HLA-DR-binding epitope (PADRE) on I-R-F (named I-P-R-F) to enhance T cell help^19^. I-P-R-F, intramuscularly injected in the lateral thigh, efficiently provides complete protection in both upper and lower respiratory tracts against a high titer SARS-CoV-2 challenge in rhesus macaques. Therefore, the IFN-armed RBD-dimer fusion proteins can be the potent COVID-19 vaccine candidates. This strategy could also be expanded to various infectious diseases, serving as a promising technology platform for vaccine development.

## Results

### I-R-F initiates highly efficient antigen presentation and shapes a favorable antibody generation environment

RBD of the SARS-CoV-2 is the primary viral protein domain to initiate cell entry and the major target for neutralization antibodies. Similar to a previous study^5^, we found that RBD presented weak immunogenicity and only induced very low titers of anti-RBD antibody even with alum adjuvant. The poor immunogenicity of RBD could be attributed to its small molecular size for antigen presentation by APCs, lacking epitopes for T cell help, and instability in vivo. It is known that organized lymphoid nodes are essential sites for better antigen presentation and interaction among DC and various lymphoid cells^20, 21^. RBD may be too small to enter lymph vessels effectively before diffusing to the surrounding muscular tissues. We, therefore, constructed an RBD-Fc fusion protein, a dimerized RBD via Fc of human Ig, which can increase protein stability and size for effective LN-targeting and FcR^+^ DC capture. To further increase antigen processing and presentation by DC, we armed RBD with type I IFN by fusing mouse IFNα at the N-terminus of the RBD-Fc to form a natural dimer, named I-R-F (Figure. 1a). I-R-F was well expressed in 293F cells and easily purified from the supernatant with Protein-A Sepharose column as previously described^22^. After a one-step Protein-A column purification, a single peak of intact I-R-F fusion protein was obtained using size exclusion chromatography. Additional SDS-PAGE also confirmed the right size of the purified fusion protein (Fig. 1b). Real-time binding kinetics showed a high binding affinity of I-R-F to hACE2 (K_D_=10.8 nM, determined by the BIAcore T100 system), suggesting that RBD in the I-R-F fusion protein was efficiently exposed (Fig. 1c). To further evaluate the antigen epitope exposure of RBD in I-R-F fusion protein, we used three RBD-specific MAbs (BD-368-2, BD-604, and BD-623) to detect the epitopes corresponding to the antibodies. BD-368-2 BD-604 and BD-623 were all isolated from SARS-CoV-2 convalescent patients and specific recognition of SARS-CoV-2 RBD. BD-368-2 recognizes the epitope in the far concern of RBD regardless of the spatial conformations^2, 11^. BD-604 belongs to the class 1 Nabs with a similar RBD-binding pose, which binds to epitopes overlapping with the RBD-ACE2 binding interface and binds only to up RBDs, not the down RBDs^2, 23^. BD-623 is Class 2 Nabs with a long CDRH3, which recognize the epitopes overlap with the ACE2-binding site and can bind to the up and down RBDs^23, 24^. The result showed all three MAbs bound to the I-R-F with high affinity, indicating the sufficient exposure of RBD epitopes on the I-R-F vaccine molecule (fig 1d). The IFN bioactivity of I-R-F was examined by an anti-viral infection assay. I-R-F could inhibit VSV-GFP infection in L929 cells in a dose-dependent manner (Fig. 1e). The function of the Fc domain was determined by RAW264.7 cell binding assays, showing the binding capacity of I-R-F to APCs (Fig. 1f).

**Figure 1.**
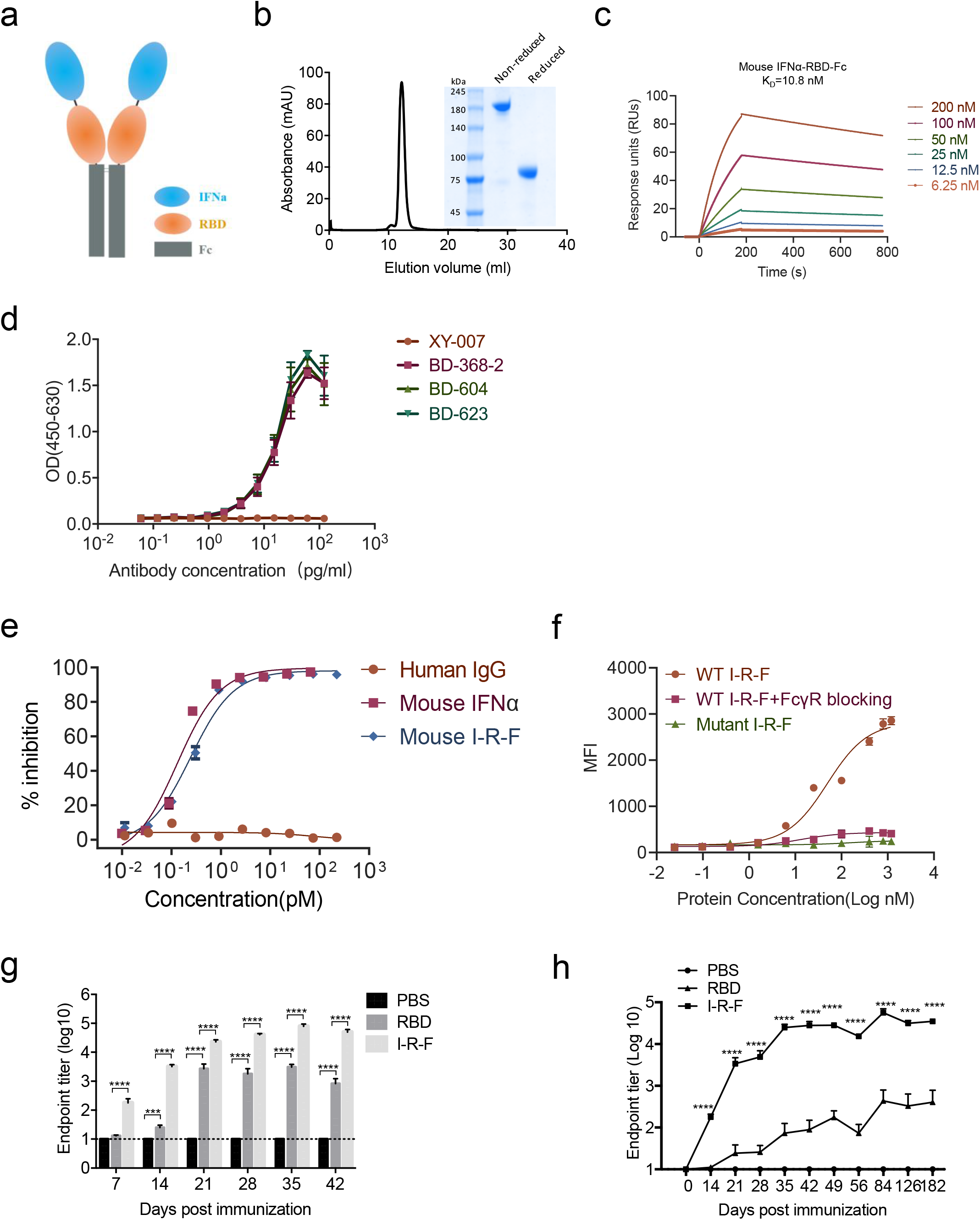
IFN-armed I-R-F induces robust IgG response. **(a)** Schematic illustration of the I-R-F fusion protein. The structural elements contain a mouse IFNα4a (IFNα), receptor-binding domain (RBD), immunoglobulin Fc fragment (Fc). **(b)** Size exclusion chromatography of I-R-F was performed on a Superdex200 Increase Column. The ultraviolet absorption at 280mm is shown. The insert photograph presents the SDS-PAGE of the eluted protein samples. **(c)** The real-time binding kinetics between I-R-F and hACE2 was determined by the BIAcore T100 system. (d) Evaluation of the binding ability of monoclonal antibodies to I-R-F by ELISA. To determine the known RBD epitopes in I-F-R, ELISA Plate was coated with I-R-F. Then, 2-fold serially diluted monoclonal antibodies were detected. The anti-Pres1 XY007 monoclonal antibody was given as the control. The absorbance was read at 450-630. (e)The bioactivity of IFNα contained in I-R-F was measured by an anti-viral infection biological assay. **(f)** Binding of mouse IFNα-RBD-Fc to FcγR on RAW264.7 cells. Cells were incubated with serial dilutions of WT IgG fusion protein, mutant IgG fusion protein, or WT IgG fusion protein with anti-FcγR, followed by a fluorophore-conjugated anti-human IgG secondary antibody. Flow cytometry measured MFI (n=3). **(g)** BALB/c mice (n=8/group) were immunized intramuscularly with 10μg of I-R-F or equimolar RBD protein, mixed with alum adjuvant, respectively. Mice were re-immunized with the same dose of vaccine on day 14 post the first shot. PBS containing alum adjuvant was chosen as a negative control. Sera were collected on days 7, 14, 21, 28, 35, and 42 after initial immunization, and the IgG levels were measured by ELISA. **(h)** Groups of BALB/c mice (n=7/groups) were intramuscularly immunized twice on day 0 and day 14 with 10μg of alum-adjuvanted I-R-F, or equimolar RBD or alum alone as a control. Serum was collected at indicated time points, and the kinetics of the RBD-specific IgG antibody titers were determined by ELISA. The dashed line indicates the limit of detection. The data shown are presented as mean ± SEM. P-values were determined by one-way ANOVA with multiple comparison tests. ns (not significant), *P<0.05, **P<0.01, ***P<0.001, ****P<0.0001.

To compare the immunogenicity of RBD and I-R-F, mice were vaccinated with 10 μg I-R-F or equimolar RBD protein formulated with alum adjuvant using a prime-boost vaccination schedule on days 0 and 14, respectively. We observed a much higher RBD-specific IgG response in I-R-F vaccinated group than in RBD vaccinated group. The immune responses were induced quickly in I-R-F groups, and the viral-specific IgG could be detected as early as 7 days after immunization. In contrast, the sera collected from RBD groups presented a much weaker and more delayed antibody response (Fig. 1g). The long-lasting and strong antibody response has remained more than 6 months (Fig. 1h). Moreover, the equivalent molar of I-R-F induced stronger humoral immunity than RBD-Dimer and RBD-Fc (Extended Data Fig. 1a and b). IFNα has been widely used in clinical treatment, and it can also be used as an adjuvant for vaccines ^25^. To determine whether the use of IFNα as an adjuvant instead of I-R-F could also induce a robust immune response, a mixture of RBD-Fc plus IFNα was compared with I-R-F. As shown in extended Data Fig. 1a, better humoral immunity was induced in the mixture group than alone R-F, yet not equal the I-R-F. Then, mice were immunized twice either intramuscularly or subcutaneously with 10 μg of alum-adjuvanted I-R-F. We found that both intramuscular and subcutaneous immunizations generated strong and long-lasting IgG responses (Extended Data Fig. 1c). Notably, the adverse effects of vaccination could not be ignored. There were no obvious body weight changes observed even in mice immunized with an extremely high dose of 100 μg I-R-F (Extended Data Fig. 6a). Moreover, only IL-6 slightly increased among mouse inflammatory cytokines (IL-12, IFN-γ, IL-6, TNF-α, and IL-10) in mice immunized with 50 or 100 μg I-R-F (Extended Data Fig. 6b). Neither ALT nor AST elevated in all groups (Extended Data Fig. 6c). Further, no IFNα specific antibody was detected in mice immunized with I-R-F (Extended Data Fig. 6d). Therefore, I-R-F overcomes poor immunogenicity of monomeric RBD, further improves the immunogenicity of RBD-Dimer and RBD-Fc, and induces long-lasting neutralizing antibodies without side effects.

### A single or low dose of I-R-F induces robust neutralization antibodies even without additional adjuvant

To determine the appropriate doses for long-lasting antibody production, mice were intramuscularly immunized with variant doses of alum-adjuvanted I-R-F (from 10 μg to 0.001 μg) twice on days 0 and 14, respectively. The level of viral-specific IgG induced by I-R-F was in a dose-dependent manner. Impressively, I-R-F, even at a low dose of 0.01 μg, could generate strong and long-lasting IgG responses (Fig. 2a). Sera from immunized mice were collected to determine the neutralizing antibody titers with live SARS-CoV-2 by focus-reduction neutralization test (FRNT). I-R-F group presented a much higher titer of neutralizing antibody than the RBD group. Even if mice were immunized with a high dose of RBD (10 μg) twice, they only generated moderate levels of antibody to RBD (Fig. 1g), which failed to block virus infection *in vitro* (Fig. 2b). These results suggest that some IgG detected in RBD vaccinated mice was non-neutralizing antibody, and a certain level of antibody binding to RBD does not automatically convert to viral neutralization. The I-R-F vaccinated mice generated higher neutralizing-antibody titers than most convalescent COVID-19 patients, especially those with mild to moderate symptoms. Some patients with mild symptoms even had no detectable neutralizing antibody (Fig. 2c). To further evaluate I-R-F’s potency, mice were intramuscularly (i.m.) immunized with only one dose of alum-adjuvanted I-R-F (10 μg) or equal molar of RBD. A strong and durable anti-RBD-specific IgG was detected only in I-R-F vaccinated mice but not in the RBD group (Fig. 2d). We further compared traditional double- or triple-dose immunization with one-dose administration in extend Data Fig. 1d. Impressively, a single dose of I-R-F induced strong and long-lasting IgG responses, similar to repeated immunization (Extended Data Fig. 1d).

**Figure 2.**
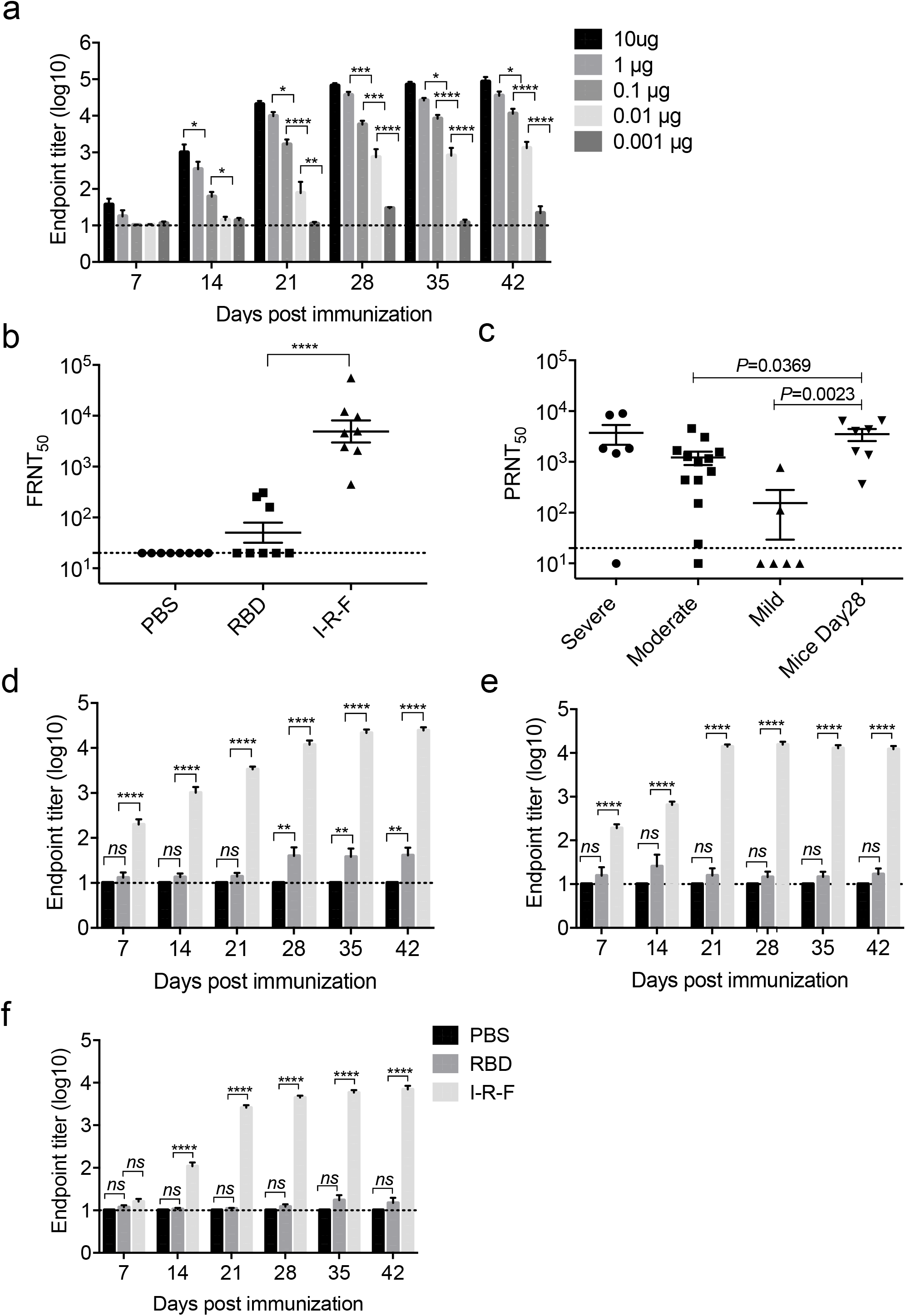
I-R-F induces robust neutralization antibodies with extreme low-dose, single-dose, and without adjuvant. **(a)** The kinetics of the RBD-specific IgG antibody response. BALB/c mice (n=7/group) were immunized with a 1:10 series dilution of the vaccine, containing 10μg, 1μg, 0.1μg, 0.01μg, 0.001μg of I-R-F, respectively. Sera were collected to assess the levels of RBD-specific IgG. **(b)** The serum described in Fig 1e on day 28 was used to determine the neutralization activity with live SARS-CoV-2 by FRNT. The sera from different groups were serially diluted and mixed with 600 FFU of SARS-CoV-2, and the mixtures were then transferred to Vero E6 cells. The number of SARS-CoV-2 foci was counted in the next day. The FRNT_50_ was defined as the reciprocal of serum dilution, which inhibits 50% of viral infection. **(c)** Comparison of the neutralizing antibody titers in sera between the I-R-F vaccinated mice (n=7) and the convalescent COVID-19 patients with different severity (n=6-13/group). The sera from 3 groups of COVID-19 convalescent patients and the I-P-F immunized mice as mentioned above in Fig 1f were serially diluted and mixed with live SARS-CoV-2. The mixtures were added to Vero E6 cells. The neutralizing titers were presented by FRNT_50_, which was the reciprocal of serum dilution neutralizing 50% of viral infection. **(d)** Mouse RBD-specific IgG induced by single immunization. Mice (n=8/group) were i.m. immunized once with alum-adjuvanted 10ug I-R-F (n=8), equimolar RBD protein, or alum alone, respectively. The levels of RBD-specific IgG in sera on days 7, 14, 21, 28, 35, and 42 after the first immunization were determined by ELISA. **(e)** Mouse RBD-specific IgG induced by vaccines without adjuvant. Mice (n=7 or 8) were vaccinated with no adjuvant 10μg I-R-F, equimolar RBD protein, or PBS control and boosted with the same dose 14-day after the initial immunization. Sera were collected every week after immunization and used to determine the IgG titers. **(f)** BALB/C mice (n=8/group) were immunized i.m. with alum-adjuvanted 0.1μg I-R-F, equimolar RBD protein, or alum alone, respectively, and boosted with the same dose at a 14-day interval. Sera were collected on days 7, 14, 21, 28, 35, and 42 after the initial immunization. RBD-specific IgG levels were analyzed by ELISA. The dashed line indicates the limit of detection. Data are shown as mean ± SEM. P-values were calculated by one-way ANOVA with multiple comparisons tests. ns (not significant), *P<0.05, **P<0.01, ***P<0.001, ****P<0.0001.

To determine if IFNα can enhance the RBD immunogenicity in the absence of adjuvant, the mice were i.m. immunized with 10 μg of I-R-F or equimolar RBD protein without alum. Again, a robust and long-lasting RBD-specific IgG was detected merely in I-R-F without adjuvant but not in the RBD group (Fig. 2e), indicating that IFNα in the fusion protein functions as a natural adjuvant to enhance the vaccine-induced immune responses. Furthermore, mice were i.m. immunized twice (day 0/14) with low dose (0.1 μg) alum-adjuvanted I-R-F or equimolar RBD, respectively. RBD group failed to produce detectable RBD-specific IgG while the I-R-F group produced high level and long-lasting antibodies to RBD (Fig. 2f). Impressively, we observed that a single dose of I-R-F, even without adjuvant or immunized at the low dose, could also generate high titers of neutralizing antibody, as shown in the pseudovirus neutralization assay (Extended Data Fig. 1e). These results demonstrate that I-R-F could be a potent anti-virus vaccine even under disadvantaged conditions, such as a single injection, with no adjuvant, or containing a trace amount of antigen.

### I-R-F induces SARS-CoV-2 specific B cell, Th1, Th2, and CD8^+^ T cell immune responses

Protein antigens with alum-adjuvant often favor Th2-biased antibody responses instead of strong Th1 and CD8^+^ T cell responses ^26, 27^, which are the crucial anti-viral response arms. To determine the antigen-specific memory B cells, we performed a B-cell ELISPOT assay three months after immunization and observed a significantly higher number of RBD-specific B cells in the I-R-F group than the RBD group, (Fig 3a and 3b, extended Data Fig. 2a and 2b). IgG1 is associated with Th2 responses in mice, and IgG2a commonly indicates Th1 responses ^28, 29^. To determine which type of immune responses were induced by I-R-F, mice were immunized with 10μg of I-R-F or equimolar RBD in alum, and the different subtypes of IgG were determined by ELISA. I-R-F induced not only IgG1 (Th2) responses but also IgG2a (Th1), while RBD only raised IgG1 responses (Fig. 3c). To further confirm the subtype of T cell responses, splenocytes from immunized mice were collected and stimulated with a peptide pool of SARS-CoV-2 RBD. Both CD4^+^ and CD8^+^ T cells and IFN-γ and IL-4 secreting cells were determined. I-R-F groups produced strong Th1 and Th2 responses while RBD groups only caused weak IL-4 responses but not IFN-γ responses at all (Fig. 3d and 3e, extended Data Fig. 2c and 2d).

**Figure 3.**
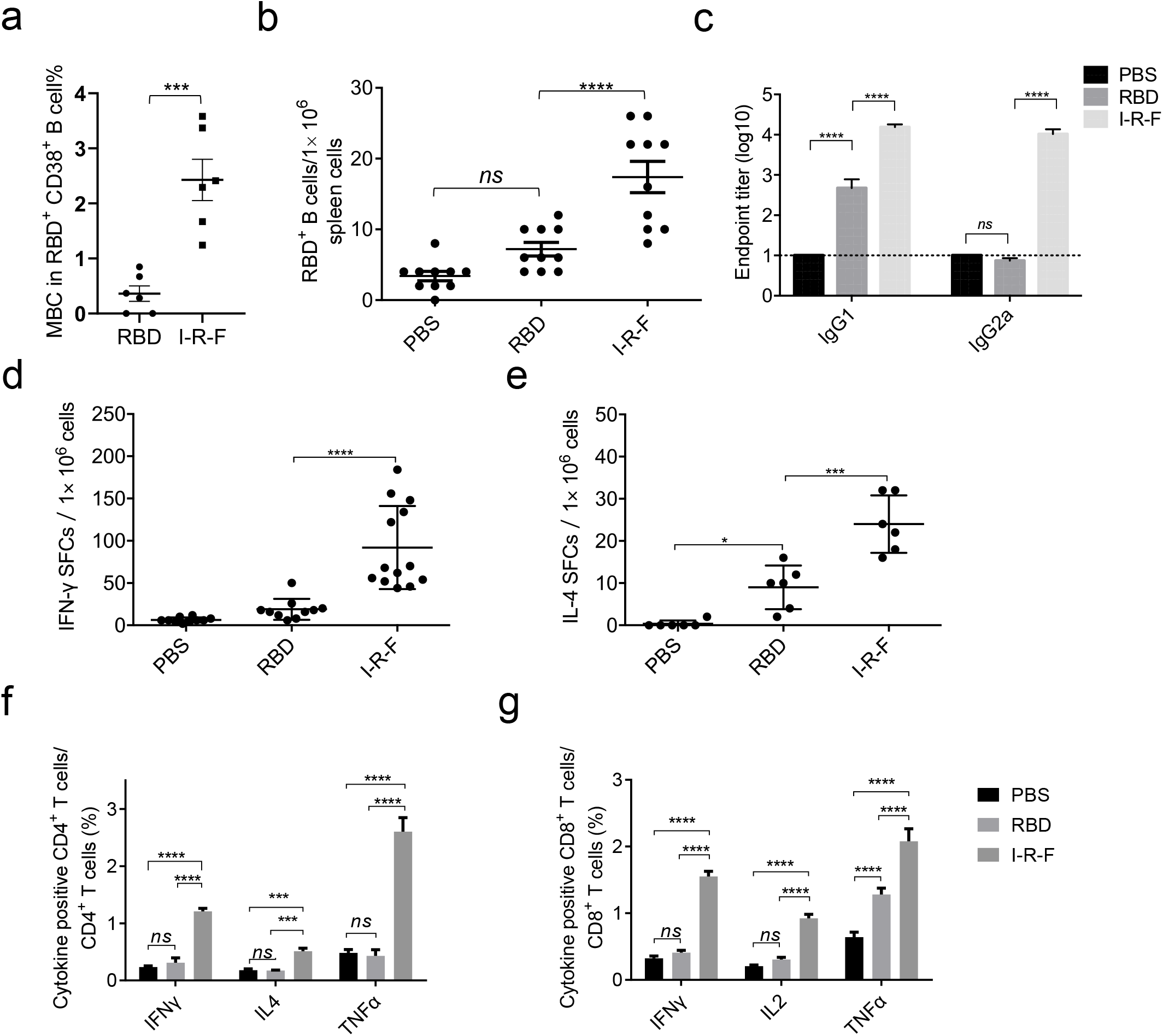
I-R-F induces SARS-CoV-2 specific Th1, Th2, and CD8^+^ T cell immune responses. (a and b) C57BL/6 mice were immunized with 10 μg I-R-F, equimolar RBD, or PBS with a prime-boost vaccination regimen in a 14-day interval. Mice were sacrificed six months post the first vaccination, and splenocytes were collected. (a) ELISPOT assay was performed to determine RBD-specific B cells in the spleen. (b) The percentage of memory B cells in the spleen was analyzed. (c) BALB/c mice (n=8/group) were immunized with 10 μg of I-R-F, equimolar RBD, or PBS twice in a 14-day interval. PBS was performed as a negative control. Sera were collected on day 28 after the initial immunization and used to determine the IgG subclasses. (d-g) C57BL/6 mice were immunized with 10 μg I-R-F, equimolar RBD, or PBS with a prime-boost vaccination regimen in a 14-day interval. Mice were sacrificed, and splenocytes were collected 28 days post the first vaccination. ELISPOT assay was performed for IFN-γ (d) and IL-4 (e) secretion from mice splenocytes stimulated with RBD peptide pool. Splenocytes were incubated with an RBD peptide pool. (f) The percentages of IFN-γ^+^, IL-4^+^, and TNF-α^+^ CD4^+^T cells were determined by ICCS. (g) The percentages of IFN-γ^+^, IL-2^+^, and TNF-α^+^ CD8^+^T cells were determined by ICCS. The dashed line indicates the limit of detection. Data are presented as mean ± SEM. P-Values in (a-e) were calculated by one-way ANOVA with multiple comparisons tests. P-values in (f) and (g) were calculated by two-way ANOVA with multiple comparisons tests. ns (not significant), *P<0.05, **P<0.01, ***P<0.001, ****P<0.0001.

The proportions of CD4^+^ T cells producing IFN-γ, IL-4, and TNFα were determined by intracellular cytokine staining. It is shown that CD4^+^ T cells in I-R-F vaccinated groups produced all three cytokines while CD4^+^ T cells in RBD vaccinated groups produced none of these cytokines, similarly as that in the unimmunized group (Fig. 3f and extended Data Fig. 2e). To determine if CD8^+^ T cells respond to viral-specific RBD, the intracellular cytokine staining was performed to determine the proportion of IFN-γ producing CD8^+^ T cells after stimulation with the RBD peptide pool. Notably, I-R-F induced a much higher number of cytokine-producing CD8+ T cells in vaccinated mice, suggesting that poor immunogenicity of RBD may not result from lacking the proper T and B cell epitopes (Fig. 3g and extended Data Fig. 2f). Therefore, I-R-F can induce potent and comprehensive T- and B-cell responses, addressing the main challenge for protein-based COVID-19 vaccines.

### I-R-F efficiently stimulates Tfh and GC generation via targeting DC in LN

We hypothesize that the more robust immune responses induced by I-R-F compared to RBD might be attributed to the fused dimerized Fc, which improves draining to LN. To demonstrate this hypothesis, we labeled RBD and I-R-F protein with fluorescein. After subcutaneous injection at the tail base, bilateral inguinal LN were isolated at different hours post injection. A significantly increased intensity of fluorescence in the I-R-F group was observed from 6 hours to 24 hours post injection, with the maximum difference shown at 12 hours (P-value < 0.0001) (Figure. 4a and 4b). Fluorescein labeled RBD failed to reach LN effectively, likely due to its smaller molecular size. In consideration of different labeling efficiency across different proteins, which may influence the quantitative analysis of fluorescent molecules, we replaced RBD with eGFP (enhanced green fluorescent protein) to generate IFNα-eGFP-Fc fusion protein (I-E-F). In order to assure consistency with the recommended immunization pathway, we intramuscularly injected equal moles of eGFP and I-E-F in the lateral thigh of mice. We analyzed eGFP^+^ myeloid cells in isolated DLN by flow cytometry. The percentages of eGFP positive dendritic cells (DCs) (Fig. 4c and 4d) and macrophages in I-E-F vaccinated mice were remarkably higher (Extended Data Fig. 3a and extended Data Fig. 3b) than that in the eGFP alone group. These data suggest that I-R-F is likely to target LN much more efficiently than RBD. To directly examine the maturation of DCs, likely resulting from IFNα stimulation, the expression of CD80 and CD86 in the I-R-F group was determined by flow cytometry. Compared to RBD, the I-R-F group presented much higher levels of CD80 and CD86 on DCs (Fig. 4e and Extended Data Fig. 3c).

**Figure 4.**
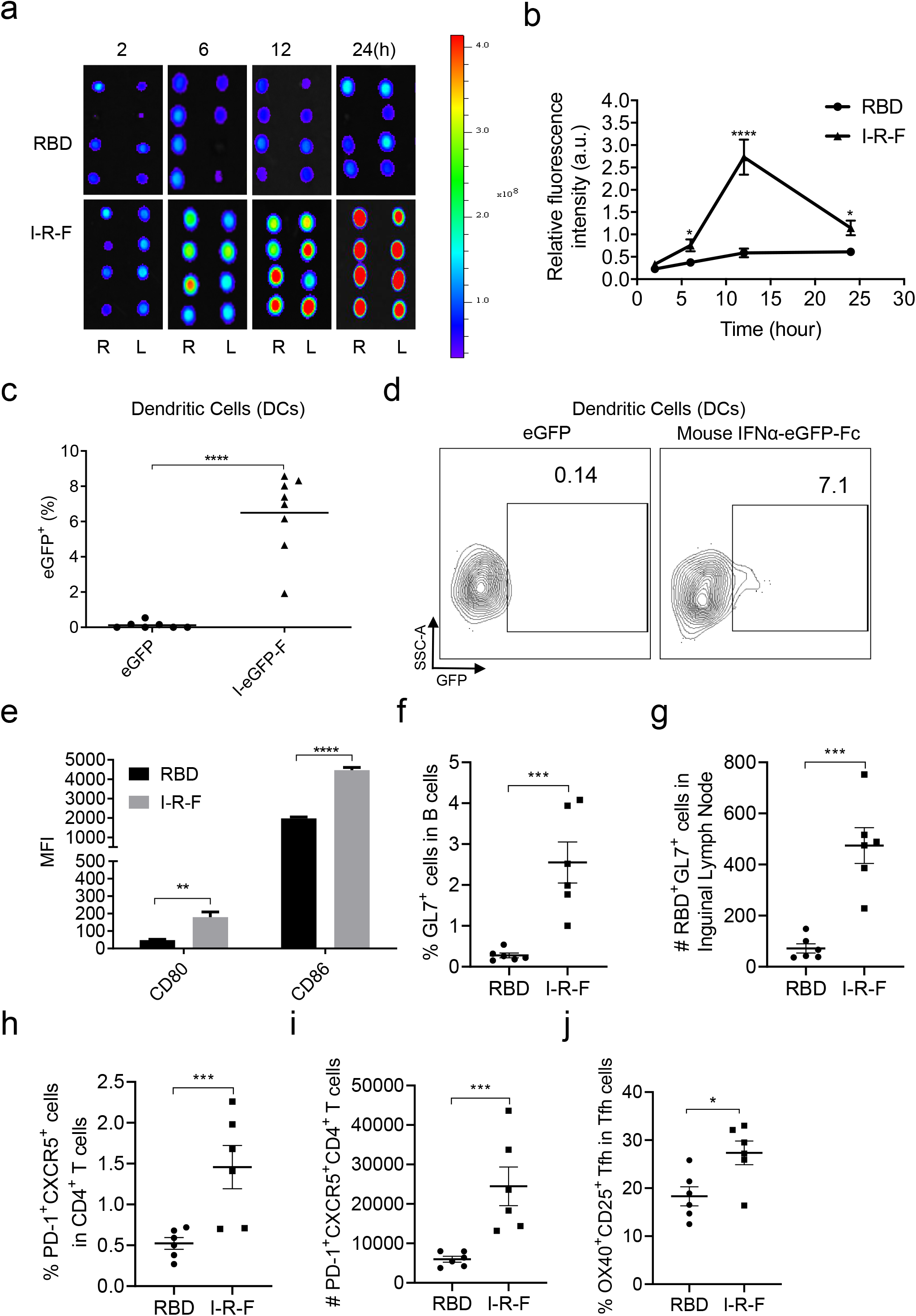
I-R-F efficiently stimulates Tfh and GC generation via targeting DC in lymph node. (a-b) BALB/c mice (n=4/group) were subcutaneously injected Cy-5.5 labeled I-R-F or RBD at the tail base, and the mice were subjected to monitor the accumulation of labeled proteins in the inguinal lymph node at the indicated time points (a) and the fluorescence intensity of the DLN was quantified using the Living Imaging software (b). (c) C57BL/6 mice (n=7 or 8/group) were intramuscularly injected 1nmole of I-E-F (mouse IFNα-eGFP-Fc) or eGFP. Four hours after injection, mice were sacrificed. Lymphocytes from mouse iLNs were collected to analyze the capture of I-E-F or eGFP. The eGFP^+^ DCs (B220^-^CD11c^hi^MHC-II^+^) by flow cytometry. The proportions of eGFP^+^ cells were analyzed. (d) The representive typical FCM figures. (e) C57BL/6 mice (n=5/group) were i.m. immunized with 10 μg of mouse IFNα-RBD-Fc (I-R-F) or equimolar RBD. Mice were sacrificed 24 hours later, DCs from iLNs were collected and analyzed with the maturation markers of CD80 and CD86 by flow cytometry. (f-j) C57BL/6 mice (n=6/group) were i.m. vaccinated with I-R-F or RBD, and inguinal LN were collected 14 days after the initial immunization. (f) The proportion of GC cells (B220^+^CD3^-^GL7^+^CD38^+/-^) was determined. (g) The number of RBD^+^GL7^+^ B cells per inguinal lymph node was counted. (h) The frequency of Tfh (CD3^+^CD4^+^PD-1^+^CXCR5^+^) was detected in each group. (i) The number of Tfh (CD3^+^CD4^+^PD-1^+^CXCR5^+^) per inguinal LN was detected in each group. (j) The Frequency of RBD-specific Tfh cells post stimulated with RBD peptide pool was measured by the proportion of CD25^+^OX40^+^ Tfh cells. Data are shown as mean ± SEM. P-values in (b-d) were analyzed with the unpaired t-test. P-values in (e-f) were calculated by one-way ANOVA with multiple comparisons tests. ns (not significant), *P<0.05, **P<0.01, ***P<0.001, ****P<0.0001.

Effective generation of T follicular helper cells (Tfh) could result from better antigen presentation to CD4^+^ T cells. Tfh in lymphatics tissues are essential for forming germinal centers (GC) and differentiation of B cells ^30, 31^. We also found the increased percentage and number of total GC B cells (Fig. 4f and extended Data Fig. 3d) and RBD-specific GC B (Fig. 4g and extended Data Fig. 3d) cells in the I-R-F group than in the RBD group (P <0.001). Also, a higher percentage and more Tfh cells were detected in inguinal LNs. Characterization of antigen-specific Tfh cells is essential to define the mechanistic basis of antibody responses. Further, an assay of activation-induced markers (AIM) was performed, showing that more RBD-specific Tfh cells were induced in I-R-F than the RBD group (Fig. 4j and extended Data Fig. 3f). Together, these data suggest that I-R-F can target and activate DC in LN more efficiently than RBD, leading to stronger Tfh and GC reactions.

### The pan DR epitope(Pan)further enhances the immunogenicity of I-R-F vaccine

To avoid the potential limitation and competition with RBD immunogenicity from dominant epitopes outside RBD in the S glycoprotein, we selected RBD-dimer as the only viral antigen. However, RBD, a smaller polypeptide portion of S protein, might contain a limited number of helper T cell epitopes to B cells and CTL in a broader population. The pan DR-binding epitope (PADRE) is known to provide broad T cell help via binding to common human HLA-DR types and mouse IA^b^ ^32, 33^. To reduce the risk of limited helper epitope for some HLA-DR, we inserted PADRE into I-R-F and constructed the I-P-R-F vaccine to augment T cell responses (the schematic model in Extended Data Fig. 4a). A clear peak of intact I-P-R-F fusion protein was visualized using size exclusion chromatography after a one-step protein-A column purification (Extended Data Fig. 4b). Additional SDS-PAGE also confirmed the purity of this fusion protein (Extended Data Fig. 4b). Real-time binding kinetics showed the high-affinity binding of I-R-F to hACE2, determined by the BIAcore T100 system (Extended Data Fig. 4c). The bioactivity of IFNα in I-P-R-F is as high as the free IFNα molecule measured by an anti-viral infection biological assay (Extended Data Fig. 4d). The mice were vaccinated i.m. with a low dose of the indicated vaccines (0.1 μg). We observed an over 10-fold increase of antibody level in mice immunized with I-P-R-F than I-R-F (Fig. 5a). The neutralization activity in the anti-sera from vaccinated mice was also evaluated using a pseudovirus neutralization assay. Consistently, mouse I-P-R-F resulted in up to 10-fold higher levels of neutralizing antibody at a notably low dose (Fig. 5b). We also generated a human I-R-F and a human I-P-R-F with a site-mutation (Q124R) on IFNα featuring an increased binding affinity of human IFNα to mouse IFNα receptor and thus allow this human IFN to be functional in the mouse model ^34^. Higher antibody titers in immunized mouse sera were induced by human IFNα-RBD-Fc than RBD-Fc, confirming the potency of human IFN in the mouse model. Furthermore, human IFNα-Pan-RBD-Fc triggered a much higher antibody response to RBD than human IFNα-RBD-Fc, which confirmed the role of Pan in enhancing the immunogenicity of poor antigen (Fig. 5c).

**Figure 5.**
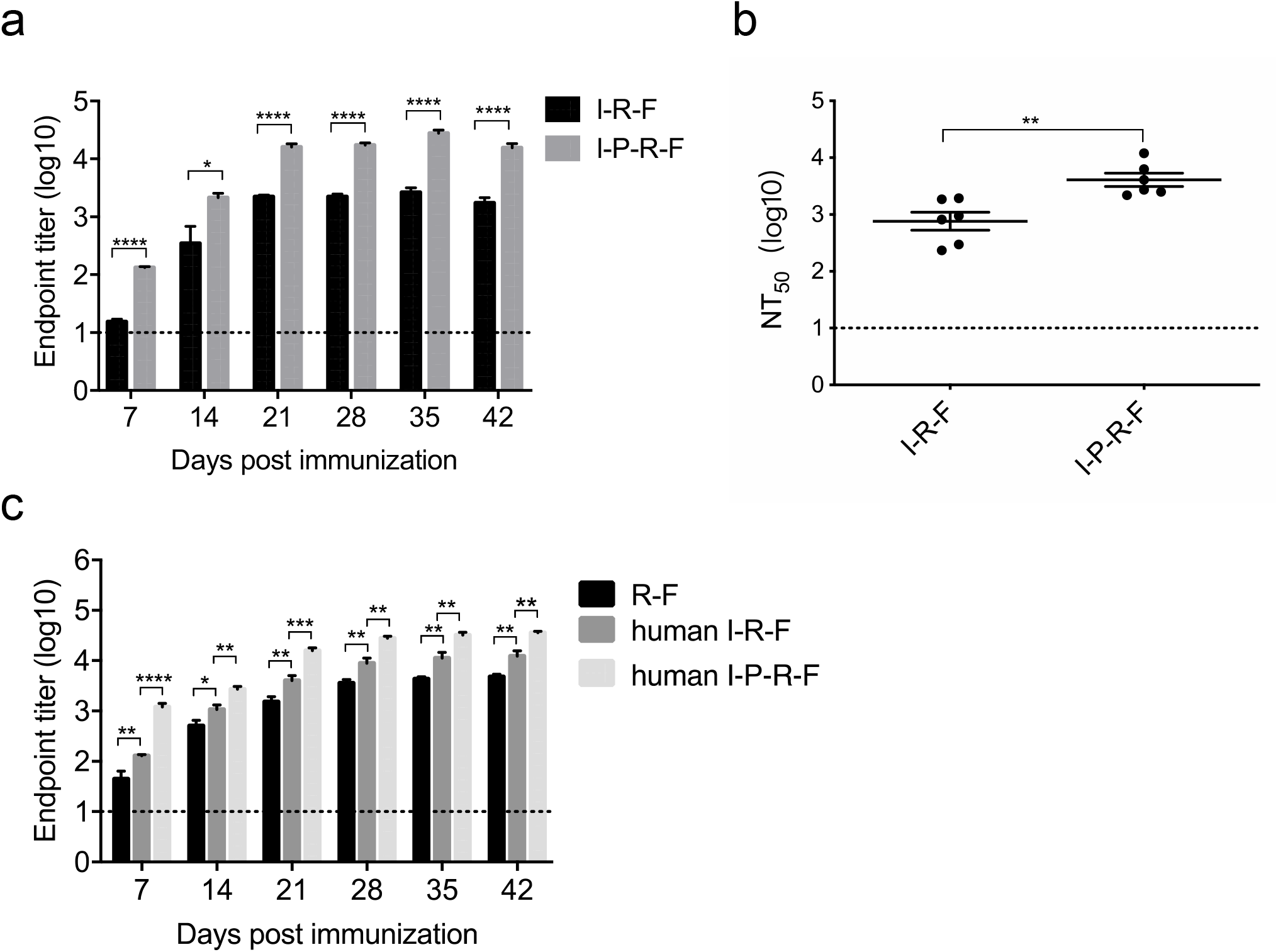
The pan DR epitope(Pan)further enhances the I-R-F immunogenicity. C57BL/6 mice (n=6/group) were vaccinated intramuscularly with 0.1μg of I-R-F or I-P-R-F using a two-dose immunization procedure. Sera were collected on days 7, 14, 21, 28, 35, 42 after the initial immunization. The RBD-specific IgG antibody titer was determined by ELISA. (b) The neutralization activity of vaccinated sera collected on day 28, as shown in (a), was evaluated using a pseudovirus neutralization assay. (c) BALB/C mice (n=8/group) were immunized i.m. with 1 μg of human IFNα-RBD-Fc (human I-R-F), human IFNα-Pan-RBD-Fc (human I-P-R-F), or equimolar RBD respectively, and boosted with the same dose at a 14-day interval. Sera were collected on days 0, 14, 21, 28, 35, and 42 after initial immunization and analyzed by ELISA. The dashed line indicates the limit of detection. Data are shown as mean ± SEM. P-values were calculated by one-way ANOVA with multiple comparisons tests in (a) and (c). P-values in (b) were analyzed with an unpaired t-test. P-values in (d-e) were determined using a paired t-test. ns (not significant), *P<0.05, **P<0.01, ***P<0.001, ****P<0.0001.

### I-P-R-F vaccination induces complete protection against high dose SARS-CoV-2 challenge in rhesus macaques

Rhesus macaque is a commonly used model for SARS-CoV-2 virology, pathology, immunology studies and screening anti-virus vaccines and medicines. To investigate the immunogenicity of this newly designed vaccine in rhesus macaques, eight macaques into four groups with a male and a female in each group and i.m. immunized twice on day 0 and day 14, with either 10 μg or 50 μg V0-1 (human I-P-R-F), mixed with or without alum-adjuvant. Impressively, all vaccinated groups generated very high levels of RBD-specific IgG antibodies, even with low-dose vaccination. High titer of antibody could be induced in macaques even by vaccine without adjuvant no matter high or low vaccine doses (Extended Data Fig. 5a and 5b). Importantly, the high titer anti-virus IgG has been maintained for 250 days so far. The sera from vaccinated animals were subjected to neutralization assays with pseudovirus and authentic SARS-CoV-2 (Extended Data Fig. 5c and 5d). The high viral neutralization titers (NT_50_>1000 and FRNT_50_>3000, respectively) indicate that this newly designed vaccine could induce strong and long-lasting protective immunity.

To further study if the vaccine has a protective effect, eighteen rhesus macaques were divided into three groups (6 per group) and vaccinated twice (day 0/14) with 10 μg or 50 μg alum-adjuvanted V0-1, or alum alone as a control, respectively. All immunization was done via the i.m. injection in the lateral thigh. Macaques were challenged with a high dose of SARS-CoV-2 (1×10^7^ TCID_50_) intranasally 21 days after the initial immunization. We observed very high titers of antibody against RBD in both low- and high-dose vaccinated groups as determined by ELISA (Fig. 6a). It is noteworthy that high titers of neutralizing antibody were presented in both low- and high-dose vaccinated groups as determined by pseudovirus and live virus neutralization assays (Fig. 6b and Extended Data Fig. 5e). Although disease symptoms for SARS-CoV2 infection are rather mild in Rhesus macaques, when monkeys were infected with a high dose of viruses, body temperature was higher in the control group (Fig. 6c), as measured every two days after virus challenge. Similarly, lower body weight was observed in the control group but not observed in the immunized group at all (Fig. 6d).

**Figure 6.**
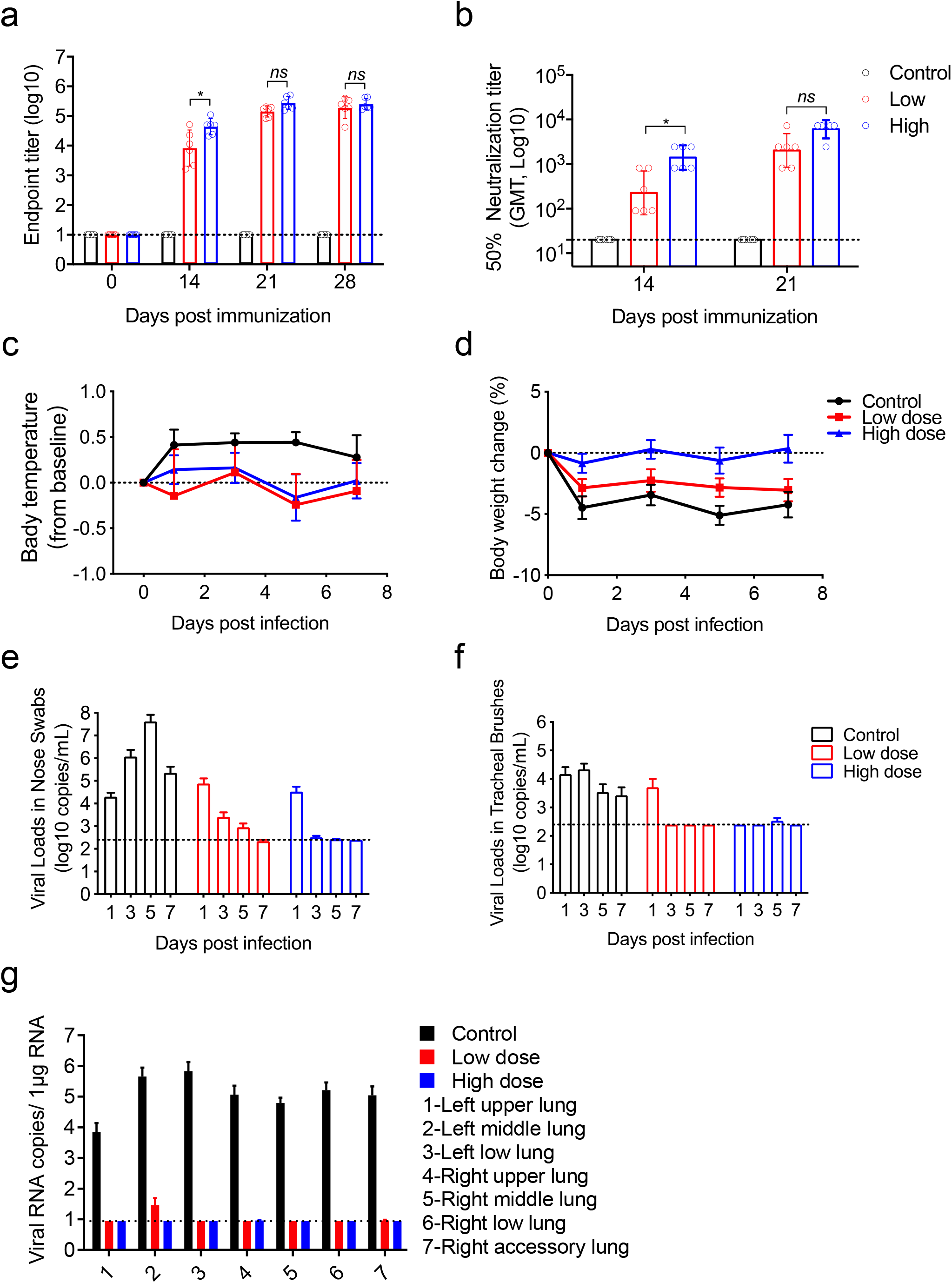
I-P-R-F vaccination induces complete protection against a high titer SARS-CoV-2 challenge in rhesus macaques. Rhesus macaques (n=6/group) were administered twice with 10μg or 50μg of alum-adjuvanted I-P-R-F or alum alone as control via the intramuscular injection and challenged with SARS-CoV-2 intranasally on day 21 after the initial immunization. Sera were collected on days 0, 14, 21 (before virus infection), and day 28 (after infection) and subjected to antibody assays. (a) The SARS-CoV-2 specific IgG was determined by ELISA. **(b)** Neutralization antibody titers were analyzed. Body temperature **(c)** and body weight **(d)** were measured after the virus challenge. Viral loads in nose swabs **(e)** and tracheal brushes **(f)** following the virus challenge were determined by qPCR. **(g)** Viral load in various lung lobes of macaques challenged with SARS-CoV-2 on day 7 post infection. Dashed line in(a-b, e-g)indicates the limit of detection. Data are shown as mean ± SEM. P-values were calculated by one-way ANOVA with multiple comparisons tests. ns (not significant), *P<0.05, **P<0.01, ***P<0.001, ****P<0.000

Virus clearance in the upper respiratory tract in rhesus macaques has rarely been reported during the first seven days of infection, even after potent vaccination. However, our vaccine resulted in a significant reduction of viral loads in the nasal passages of macaques in the low dose-vaccine group and even undetectable viral load in the high dose-vaccine group only one day after high titer virus infection, as determined by sensitive qPCR (Fig. 6e). Similarly, viral loads in tracheal brushes were also undetected in both the low- and high-dose vaccinated groups one day after virus infection (Fig. 6f). Viral loads in anal swabs of rhesus macaques were undetectable one day after infection in both the low- and high-dose vaccinated groups (Extended Data Fig. 5f). For each group of vaccinated macaques, 84 specimens from lung lobes (14 samples/7 lobes/mice) were collected on day 7 post infection and subjected to determine the viral loads in all areas of lung lobes by q-PCR. Over an average of 4 log reduction of virus load was observed in the lungs of both low- and high-dose of vaccinated groups, while extremely high viral loads were maintained in the lungs of control macaques (Fig. 6g and Extended Data Fig. 5g). These data demonstrate that V0-1 can generate highly effective protection against SARS-CoV-2 infection in both upper and lower respiratory tracts even with low-dose vaccination.

A Randomized, double-blind, placebo-controlled phase I clinical trial was initiated on Feb 24, 2021 to evaluate the safety and immunogenicity of recombinant SARS-Cov-2 fusion protein vaccine (V-01) in healthy subjects (Phase I: ChiCTC2100045108; phase II: ChiCTR2100045107). The study was a single-center clinical trial, in which healthy subjects aged 18 and above were divided into 120 for treatment while 30 for placebo) were given two doses of recombinant SARS-CoV-2 fusion protein vaccine (V-01) or placebo on days 0 and 21, respectively. All the AEs reported in each group within seven days after dosing of 10, 25, and 50 μg. The first-dose vaccination, safety follow-up on day 7 and a follow-up in all adult subjects were completed. All AEs of grade 1 (23.33%) were observed within seven days after dosing, and all the remaining AEs were of grade 2 (3.33%). There were no AEs or SAEs of grade ≥3 that were related to the investigational vaccine. All adults from 18-60 had positive for antibody to RBD. The observation is still ongoing.

## Discussion

Multiple types of SARS-CoV-2 vaccines have been developed and entered into clinical trials in an endless stream. Immunological effects and safety of these vaccines have been reported ^16, 17, 42-44^. An ideal vaccine should have the following properties: 1) efficacy and safety, presented by high titer neutralizing antibodies and T cell responses with no toxicity and antibody-dependent enhancement (ADE); 2) prolonged protective immunity and easy administration; 3) “simplified” large-scale production, storage, and distribution. Here, we designed and evaluated a vaccine platform based on IFN-α armed RBD, a dimerized human IgG1 Fc (I-R-F) molecule. Evidenced by the results on mice and rhesus macaques, the I-R-F vaccine may provide an effective prophylactic solution for COVID-19. Because 1) Potency: Due to its size, IFN-armed RBD dimerized by Fc will primarily target to DLN and increase antigen processing and presentation inside DLN. Therefore, hundreds to thousands fold more neutralizing RBD-specific IgG1 and IgG2a antibodies were induced by I-R-F than monomer RBD. In addition, Th1, Th2, and CD8^+^ T cell responses could be readily detected after vaccination; 2) Safety: RBD is the only exogenous antigen in the vaccine; thus, all antibodies induced are targeting RBD to block viral entry. I-R-F could be elicit robust and long-lasting immune responses at a low dose (0.01 ug) or even without adjuvant. Unlike smaller molecule of free IFN (18.7), much higher molecule weight of I-R-F will allow this molecule to flow into DLN to improve antigen presentation and DC mature. Alum adjuvant used in this vaccine is well known and widely used without severe toxicity. Furthermore, alum-adjuvanted I-R-F allows slow release of I-R-F molecule into DLNs, resulting in prolonged, effective, and safe immune challenges inside DLNs. On the other hand, there were no detectable autoimmune B or T cell responses to IFN. 3) Durability:The level of neutralizing antibodies induced by I-R-F vaccine was maintained for more than 250 days in the rhesus macaque model; 4) Accessibility: fusion with Fc is easy for large-scale manufacturing and purification. High productive clones (one gram level per liter) for large scale human I-P-R-F (V-01) production of have been confirmed in GMP grade manufacturing; 5) Portability:this fusion protein is stable at 4-25°C for months, which notably allows for more accessible transport and storage for developing countries.

A potential hurdle of ADE induced by antibodies with weak neutralizing activities should be considered in developing the SARS-CoV-2 vaccine ^45^. COVID-19 patients with higher antibody titers against SARS-CoV-2 may be associated with more severe illness, though the mechanism is still not fully determined ^46^. Subunit vaccines have been proposed to focus on S or RBD protein inducing the neutralizing antibodies to avoid producing ineffective antibodies and reduce ADE. However, monomeric RBD is a weak immunogenic antigen, and multiple immunizations are needed for acquiring protective immunity ^11^. To enhance immunogenicity, either modified S or RBD, such as S-trimer and RBD-dimer proteins, were developed, which help to mimic the native 3D conformation of oligomerized S protein *in vivo* ^5, 47^, or forming protein nanoparticles by displaying the RBD ^48, 49^. However, the large-scale productions of those new adjuvants remain challenged. Additionally, RBD fused with the Fc domain could also be a better choice than simple RBD or S protein^7-9^. With equivalent molar, both I-R-F and I-P-R-F induced stronger humoral immunity than RBD-Dimer and RBD-Fc. In general, protein vaccines could not have enough protective immunogenicity without additional potent adjuvants. IFN can provide an adjuvant effect with undetectable toxicity while enhancing immunogenicity and antigen presentation.

For protein vaccines, Th1 or Th2 responses are mainly dependent on types of adjuvants. Notably, enhanced immunopathology was associated with a Th2-biased response ^10, 50-52^. The choice of adjuvant is thus a crucial point to solve the problem. Alum is a universal adjuvant accompanied with protein vaccines leading to high antibody titers, while it always induces Th2-biased response ^53^. Moreover, weak or even no CD8^+^ T cell responses could be elicited by protein-based vaccines. All these drawbacks make protein vaccines undesired for the ideal and effective vaccine candidates. Herein, we designed an I-R-F vaccine with extremely strong immunogenicity even at a minuscule dose or without adjuvant. Mechanistically, multiple factors contribute to such robust immune responses. 1) IFNα is the most potent cytokine for antigen processing and presentation of DC ^54^. With I-R-F, we have observed more efficient antigen uptake by DCs; 2) IFN-α could also be used as an adjuvant in improving the generation of Tfh ^55^. We found that the I-R-F vaccine induced higher percentages of Tfh and GC B cells, as well as both high RBD-specific IgG1 and IgG2a titers indicating strong humoral immunity. Moreover, IFN might help Th1 and CTL responses even with alum. Indeed, the I-R-F vaccine could transfer the aluminum-induced Th2-biased response to a more robust Th1-Th2 balanced response. 3) Importantly, The Pan epitope further enhances the I-R-F vaccine-induced robust CD4+ and CD8^+^ T cell responses that contribute to eliminating virus-infected cells; 4) The additional Fc can allow RBD form dimer naturally and make I-R-F to become a much bigger molecule than RBD, to get to DLNs more efficiently.

High doses and long-term use of various formats of type I interferon have been approved for several viral infections, but some patients showed various toxicities^56-58^. No significant body weight loss or abnormal ALT/AST above background was observed after vaccination for preclinical studies. There was no significantly increased cytokines induced by the vaccine at the tested doses. We have not observed any sign or symptom from I-R-F vaccinated host.

In the light of significant immunogenicity, complete protection efficiency in both upper and lower respiratory tract, and undetectable toxicity of I-P-R-F in macaques, this unique protein vaccine can be potentially used in very low dosage and merely single vaccination. Furthermore, this protein vaccine maintains its potency even without adjuvant, which may provide a good chance for intranasal vaccination in all ages, especially young children. Nasal I-R-F vaccine investigations are underway since it could trigger mucosal IgA response to prevent the virus from invading mucosa in nose and upper respiratory tract or infecting others. Armed-IFN that overcomes poor immunogenicity of some antigens can be applied broadly to prevent other infectious diseases during future pandemics. This enhanced immunogenicity by new design is not limited to protein vaccine and could also be applied to other vaccine forms, such as genetic or viral-vector-based vaccines.

## Materials and Methods

### Ethics statement

All mice involved in the experiments were approved by the Biomedical Research Ethics Committee of the Institute of Biophysics of the Chinese Academy of Sciences and were performed in compliance with the Guidelines for the Care and Use of Laboratory Animals of the Institute of Biophysics. Non-human primates, Rhesus macaques immunogenicity studies were performed in the animal facility of Guangxi Fangchenggang Biotechnology Development Co., Ltd. (GFBDCL), according to the guidelines of the Committee on Animals of GFBDCL (approval No.: SYXK2018-0004/200005). Non-human primates, Rhesus macaque, infection studies were performed in the Biosafety Level 3 (BSL-3) in the Kunming National High-Level Biosafety Research Center for Non-Human Primates, Center for Biosafety Mega-Science, KIZ, CAS, according to the guidelines of the Committee on Animals of KIZ, CAS (approval No.: IACUC20005).

### Animals

Female (6-8-week-old) BALB/c mice and C57BL/6J mice were obtained from Vital River (Beijing) and bred under specific pathogen-free (SPF) conditions in the animal facility of the Institute of Biophysics and the Institute of Microbiology, Chinese Academy of Science. Eight healthy rhesus macaques were used for vaccine immunogenicity analysis. These macaques (4 male, 4 female, between 2.5-4-year old) were purchased from Guangxi Fangchenggang Biotechnology Development Co, Ltd. and housed in a clean-level animal facility of Guangxi Fangchenggang Biotechnology Development Co., Ltd. Additional eighteen healthy rhesus macaques were used for virus infection experiment. These healthy ChRMs (male, *n*=9; female, n=9: 3-5 years old) were sourced from the Kunming Primate Research Center, Kunming Institute of Zoology (KIZ), Chinese Academy of Sciences (CAS), and housed in the Kunming National High-Level Biosafety Research Center for Non-Human Primates, Center for Biosafety Mega-Science, KIZ, CAS. All animals recruited in this study are healthy and not involved in other studies.

### Cell lines, virus, and reagents

293F cells (Gibco) were maintained in SMM-TII medium (M293TII, Sino Biological), incubated in Polycarbonate Erlenmeyer Flask under 135rmp speed in an orbital shaker and cultured in an 8% incubator at 37 °C. Vero E6 cells were obtained from ATCC, and 293-ACE2 was kindly provided by Prof. Zheng Zhang (National Clinical Research Center for Infectious Disease, Shenzhen Third People’s Hospital, Shenzhen, Guangdong, China). Cells were cultured in 5% CO_2_ and maintained in Dulbecco’s modified Eagle^’^s medium (DMEM) supplemented with 10% heat-inactivated fetal bovine serum, 100 U/ml penicillin, and 100 mg/ml streptomycin.

SARS-CoV-2 pseudovirus was produced in house as previously described^8^. Briefly, human immunodeficiency virus backbones expressing firefly luciferase (pNL43R-E-luciferase) and pcDNA3.1 (Invitrogen) expression vectors encoding the SARS-VoV-2 S protein were co-transfected into 293T cells (ATCC). Viral supernatants were collected 48 h later. Viral titers were measured as luciferase activity in relative light units (Bright-Glo Luciferase Assay Vector System, Promega Biosciences).

The SARS-Cov-2 strain (Bata/Shenzhen/SZTH-003/2020, EPI_ISL_406594 at GISAID) was obtained from a nasopharyngeal swab of an infected patient, and the virus was stock propagated in Vero-E6 cells. The SARS-CoV-2 strain 107 (NMDC000HUI) was used in rhesus monkey infection. This virus strain was obtained from Guangdong Provincial Center for Disease Control and Prevention, Guangdong Province, China. The virus was stock and amplified in Vero-E6 cells.

The ploy-histidine tagged hACE2, rabbit anti-SARS-CoV-2 nucleocapsid, and HRP-conjugated goat anti-rabbit IgG (H+L) antibody antibodies were purchased from Sino Biological lnc. (Beijing, China). The peptide pool panning the SARS-CoV-2 RBD consisting of 53 peptides (15-mer) overlapping by 11 amino acids were synthesized by China Peptides Co., Ltd (Shanghai, China).

### Protein expression and purification

The COVID 2019 vaccine protein was expressed in 293F cells, as described previously^22, 59^. The coding sequence for SARS-CoV-2 RBD spanning S protein 319-541 (GenBank: YP_009724390) was codon-optimized for mammalian cells and synthesized by GENEWIZ, China. For I-R-F expression, murine IFNα4 was fused to the N-terminus of RBD with a (G4S)_4_ linker. The IFNα-RBD sequence was then cloned into the PEE12.4 (Lonza) with a human IgG1 Fc, forming the IFNα-RBD-Fc (I-P-F) fusion protein. The plasmid was transiently transfected into 293F cells. The supernatant was collected seven days after transfection, and the protein within the supernatant was purified with a Protein A-Sepharose column (GE Healthcare) according to the manufacturer’s instruction for primary purification. The eluted protein was further purified using a Superdex 200 Increase 10/300 GL column (GE Healthcare). The purity and size of the protein were analyzed by sulphate-polyacryl-amide gel electrophoresis (SDS-PAGE). R-F, I-E-F, I-P-R-F with a CD4 helper epitope (PADER), I-R-F (human) with a human IFNα2 substituting for murine IFNα4, I-P-R-F (human) were expressed and purified with the same method as described above.

SARS-CoV-2 RBD protein (rRBD) was also expressed in 293F cells. In brief, the coding sequence for RBD with a 6 x his tag on C terminus was cloned into the pEE12.4 vector without a human IgG1 Fc. The plasmid was transiently transfected into 293F cells. The supernatant was harvest on day 7, and the protein was purified using Ni-NTA agarose beads (GE Healthcare). The protein was further purified on a Superdex 200 Increase 10/300 GL size exclusion column (GE Healthcare). SDS-PAGE was performed to determine the purity and size of the protein. The eGFP and dimer-RBD protein were expressed and purified in the same method as rRBD.

### Surface plasmon resonance (SPR) analysis

Surface plasmon resonance assays were performed by a BIAcore T100 instrument with a CM5 sensor chip (GE Healthcare). Experiments were carried out at 25°C in binding buffer (PBS, 0.05% Tween 20, pH 7.4). The CM5 sensor chip was used to capture about 100 response units (RUs) The RBD of SARS-Cov-2, RBD-Fc, Mouse IFNα-RBD-Fc, Mouse IFNα-Pan-RBD-Fc, Human IFNα-RBD-Fc and Human IFNα-Pan-RBD-Fc for 3 min respectively. A two-fold serial dilution of hACE2 (from 6.25nM to 200nM) were run across the chip surface with a flow rate of 30μl /min, and the real-time response was recorded. The resulting data were analyzed fitting to a 1:1 binding model with Biacore Evaluation Software.

### The anti-viral activity of IFNα

The IFNα bioactivity was determined by the anti-viral infection assay using the L929 fibroblast cell line sensitive to VSV infection. L929 cells was seeded in 24-well plates (4 ×10^5^ per well) and incubated for approximately 16 hours. The serial dilution of I-R-F, I-P-R-F, were incubated into the medium of L929 cells and incubated for 24 hours at 37°C with 37% CO_2_. The cells were infected with 5 MOI of VSV-GFP virus and further cultured for 30 hours on the next day. Cells were collected and fixed by 4% PFA and subjected to analysis on a FACS Fortessa flow cytometer (BD Bioscience). Cells with GFP signal positive were considered as the virus-infected cells.

### Mouse vaccination

The immunogen used to immunize mice was diluted with PBS and mixed with or without a fixed-dose (20ug per mouse) of alum adjuvant (SEVA, Germany). To make alum adsorb the immunogen efficiently, the mixture was kept rolling overnight at 4°C. Female (6-8-week-old) BALB/c or C57BL/6 mice were immunized intramuscularly or subcutaneously with different immunogens in 100μL using insulin syringes. PBS containing alum was used as control. Sera were collected at indicated time points to determine the levels of SARS-CoV-2 RBD specific IgG and neutralization antibody. The detail of mouse vaccination was described in figure legends.

### Enzyme-Linked Immunosorbent Assay (ELISA)

The 96-well plates (Conning, USA) were coated with 100ul SARS-Cov-2 RBD (1.5ug/ml) overnight at 4°C. Plates were washed with PBS and blocked with a blocking buffer (PBS containing 5% fetal bovine serum, FBS) on the next day. Immunized animal serum samples were serially diluted and added into the blocked plates, followed by incubation at 37°C for 1 hour. Plates were then washed with PBST (PBS containing 0.05% Tween 20) and incubated with goat anti-mouse IgG-HRP (1:5000, Cwbiotech) or goat anti-monkey IgG-HRP (1:10000, Invitrogen) at 37°C for 30 minutes. To detect the Ig subclasses, goat anti-mouse IgG1 (1:5000, Proteintech), goat anti-mouse IgG2a (1:5000, Proteintech) were added. Plates were washed with PBST, and HRP substrate TMB was added. The reactions were stopped by 2M sulfuric acid. The absorbance at 450-630 was read using a microplate reader (Molecular Devices). The endpoint titers were defined as the reciprocal of max serum dilution at which the absorbance was higher than 2.5-fold of the background.

### Pseudovirus neutralization assay

The pseudovirus neutralization was carried out as described previously. In brief, the pseudovirus was produced by co-transfection of the plasmid expressing firefly luciferase (pNL43R-E-luciferase) and pcDNA3.1 expressing the SARS-CoV-2 spike protein into 293T cells. After 48 hours, the viral supernatant was collected, and viral titers were determined by luciferase activity in relative light units. To evaluate the neutralization of vaccinated mice serum, 293-hACE2 cells were seeded into 96-well plates (2×10^4^ per well) and 3-fold serially diluted heat-inactivated serum samples were incubated with 100 TCID_50_ of pseudovirus for 1 hour at 37°C. Medium mixed with pseudovirus was given as control. The mixture was transferred to the 96-well plates, and the platers were continued to incubate for another 24 hours. According to the manufacture^’^s instruction, the luciferase substrate was added, and luciferase activity was determined by the Bright-Lite^™^ Luciferase Assay System (Vazyme). The 50% neutralization titer (NT_50_) was defined as the reciprocal of serum dilution at which the relative light units (RUL) were reduced by 50% compared with virus control wells.

### Focus-reduction neutralization test (FRNT)

Vero-E6 cells were seed into 96-well plates with a density of 2×10^4^ per well. Sera from immunized animals and convalescent COVID-19 patients were serially diluted and mixed with 75 μl of authentic SARS-CoV-2 (8 × 10^3^ focus-forming units (FFU)/ml)). The mixture was incubated for 1 hour at 37°C and then transferred to the 96-well plates seeded with Vero E6 cells. Plates were incubated for 1 hour at 37°C. The inoculums were removed, and the plates were overlaid with medium (100 μl DMEM containing 1.6% carboxymethylcellulose, CMC). The plates were incubated for another 24 hours at 37°C. The supernatant was removed, and cells were fixed with 4% Paraformaldehyde for 30 min. Cells were subsequently permeabilized with PBS containing 0.2% Triton X-100. After PBS washing for three times, the cells were incubated with cross-reactive rabbit anti-SARS-CoV-2 nucleocapsid IgG (Sino Biological) for 1 hour at 37°C. After incubated with the primary antibody, plates were washed three times with PBST before adding the second antibody, HRP-conjugated goat anti-rabbit IgG (Jackson ImmunoReseach). Cells were further incubated for 1 hour at 37°C. After washing, the KPL TrueBlue peroxidase substrates (Seracare Life Science) were added to the plates. The supernatant was removed, and the plates were washed three times with deionized water five minutes later. The numbers of SARS-CoV-2 foci were read using an ELISPOT reader (Cellular Technology). The FRNT_50_ was defined as the sera dilution at which neutralization antibodies inhibited 50% of the viral infection.

### Authentic SARS-CoV-2 neutralization CPE assay

Vero-E6 cells were harvested and seeded into 96-wells plates with 2×10^4^ cells per well and cultured at 37°C overnight. The Serum samples from immunized Rhesus Macaques were inactivated at 56°C for 30 minutes and serially diluted with cell culture in 3-fold steps. The diluted serum samples were mixed with a virus suspension containing 100 TCID50 authentic SARS-CoV-2 in an equal volume. After neutralization in a 37°C incubator for 1 hour, the mixtures were transferred to the 96-well plates containing Vero-E6 cells. Inoculated plates were cultured in a CO2 incubator at 37°C for 6 days. Cytopathic effect (CPE) of each well was scored, and the neutralization titer was calculated as the reciprocal of serum dilution at which neutralization antibodies inhibited 50% of viral infection.

### Enzyme-linked immunospot (ELISPOT) assay

Murine IFN*γ* and IL-4 ELISPOT assays were carried out according to the manufacture^’^s protocols for mouse IFN*γ* and IL-4 ELISPOT kit (BD Bioscience). Immunized mice splenocytes were seeded in the plates with a density of 2 × 10^5^ cells per well and incubated with the peptide pool of 15 mer peptides with 11 overlapping amino acids for SARS-CoV-2 RBD protein (5 μg/mL) in pre-coated 96-well ELISPOT plates with IFNα or IL-4, Concanavalin A (ConA, Sigma) as a positive control or medium as a negative control for 48 hours at 37°C. Then, the cells were removed, and biotinylated IFN-γ or IL-4 (BD Bioscience) was added to the plates, followed by incubation for 2 hours at room temperature. The plates were washed three times with PBST before adding Streptavidin-HRP (BD Bioscience). The BD ELISPOT AEC substrate (BD Bioscience) was used to develop the spots. Spots were counted and analyzed using an automated ELOSPOT reader (Cellular Technology). For B cell ELISPOT assay, splenocytes from immunized mice were seeded in the plate coating with 2.5 μg/mL RBD protein. Cells were removed 16 hours post incubation, and biotinylated goat anti-mouse IgG (BBI life sciences) was added into plates, followed by incubation for 2 hours at room temperature. The plates were washed three times with PBST before adding Streptavidin-HRP (BD Bioscience). The BD ELISPOT AEC substrate (BD Bioscience) was used to develop the spots. Spots were counted and analyzed using an automated ELOSPOT reader (Cellular Technology).

### Intracellular cytokine staining (ICCS)

To evaluate the cytokine expression in antigen-specific T cells, Intracellular cytokine staining (ICCS) assay was performed. Mouse splenocytes were seeded in U-bottom 96-well plates (1×106/well) and stimulated with a peptide pool for SARS-CoV-2 RBD protein as described above. After 12-hour stimulation, Brefeldin A (Biolegend) was added, and the cells were collected, and first stained with anti-CD3 (Biolegend), anti-CD4 (Biolegend), anti-CD8 (Biolegend) and the LIVE/DEADTM dyes before fixed and permeabilized using a fixation/permeabilization kit (BD bioscience). The cells were subsequently performed with intracellular staining for IFN-γ (XMG1.2), IL-2 (JES6-5H4), TNF-α (MP6XT22),IL-4 (11B11). The cells were acquired using a FACS Fortessa flow cytometer (BD Bioscience) and analyzed with Flowjo software (TreeStar).

### *In vivo* imaging analysis

BALB/c mice were subcutaneously injected with 0.1 μmol (cy5.5) of different cy5.5-labeled proteins (RBD, I-R-F) at the tail base. The inguinal LNs were extracted for imaging DLN and were imaged at indicated time points post injection using an *in vivo* imaging system (IVIS) Spectrum (PerkinElmer). The fluorescence imaging data were analyzed by Living Image software (PerkinElmer).

### Flow cytometry analysis

The DLNs were excised and digested into a single-cell suspension. 2×10^6^ cells were blocked with anti-CD16/32 (anti-FcγIII/II receptor, clone 2.4G2) and stained with specific fluorescence-labeled antibodies. For evaluating antigen-presenting cell maturation in immunized mice, DCs were incubated with anti-CD11c (N418) and anti-MHCII (M5/114.15.2). For phenotypic maturation analysis, anti-CD80 (16-10A1) and anti-CD86 (GL-1) were used. For T follicular helper cells (Tfh) and germinal center (GC) B analysis, cells in DLN were stained with anti-CD3 (17A2), anti-CD4 (RM4-5), anti-CD8 (53-6.7) anti-PD1 (RMP-30), anti-CXCR5 (L138D7), anti-B220 (RA3-6B2) For *in vivo* uptake assay, the cells were labeled with anti-CD11c (N418), anti-MHCII (M5/114.15.2), anti-CD11b (M1/70) and anti-F4/80 (BM8). All the samples were acquired by BD LSRFortessa flow cytometer (BD Bioscience), and the data were analyzed with Flowjo software (TreeStar).

### Immunogenicity of I-P-R-F in rhesus macaques

To evaluate the immunogenicity of I-P-R-F in non-human primates, a total of 10 rhesus macaques (5 male and 5 female, weighing 3∼5 kg) purchased from Guangxi Fangchenggang Biotechnology Development Co., Ltd. were randomly assigned into four groups with one male and one female in each group and intramuscularly immunized with high does(50 μg) or low dose(10 μg) of I-P-R-F with or without alum as adjuvant two times at a 14-day interval. Blood was collected at indicated time points, and the SARS-CoV-2 specific IgG and neutralization antibody titers in serum were determined.

### SARS-CoV-2 challenge in rhesus macaques

A total of 18 Rhesus macaques (3-5-year old) were recruited and assigned into three groups. Macaques in group 1 were intramuscularly immunized with PBS formulated with the alum-adjuvant as control. Group 2 and Group 3 were vaccinated with 50 μg (high dose) or 10 μg (low dose) of I-P-R-F. Animals received a two-dose immunization procedure on day 0 and day 14 after the initial vaccination. The macaques were intranasally challenged with 10^7^ TCID_50_ of SARS-CoV-2 seven days after the second immunization. Animal body temperature and body weight were recorded after the virus challenge. Serum was collected on days 0, 14, 21, and 28 after challenge and subjected to antibody assays. The viral loads in nose swabs, tracheal brushes, and anal swabs were determined by qRT-PCR at indicated time points, and the lung tissues were collected on day 7 post infection and used to determine the viral load.

### Quantitative reverse transcription-polymerase chain reaction (qRT-PCR)

Viral RNA in swabs, brushes, and lung tissues was determined by quantitative reverse transcription PCR (qRT-PCR). In brief, total RNA in swabs and brushes were extracted using the QIAamp viral RNA Mini kit (QIAGEN) according to the manufacturer^’^s instruction. Lung tissues were homogenized, and RNA was extracted with an RNeasy Mini kit (QIAGEN). The viral RNA copies were determined using THUNDERBIRD^™^ probe one-step qRT-PCR kit (TOYOBO) with the following primers and probes: forward primer 5’-GGGGAACTTCTCCTGCTA GAAT-3’, reverse primer 5’-CAGACATTTTGCTCTCAAGCTG-3’, and probe FAM-TTGCTGCTGCTTGACAGATT-TAMRA-3’. SARS-CoV-2 RNA reference standard (National Institute of metrology, china) was serial diluted and performed to generate the standard curve.

### Quantification and statistical analysis

All statistical analyses were performed using Graphpad Prism 8.0. Data are shown as the mean ± SEM. An unpaired student’s two-tailed *t-test* was used to determine statistical significance for comparison between two groups. One-way ANOVA with Turkey’s multiple comparison test or two-way ANOVA with Turkey’s multiple comparisons test was conducted to compare differences among multiple groups. P values of < 0.05 were considered significant. p<0.05 (*), p<0.01 (**), p<0.001 (***) and p<0.0001 (****). *ns*, no significance.

## Acknowledgements

We are grateful to Professor Xioaliang Sunney Xie for providing monoclonal antibodies to RBD, Professor Xiaoli Wang and Dr. Xiaohan Yin for statistical analysis, Kaiting Yang for the graphic draw, and Sherry Parker and May Lynne Fu for their comments on the project.

We appreciate the funding from the National Key R&D Program of China (2018ZX10301-404) to Hua Peng, Emergency Key Program of Guangzhou Laboratory, Grant No. EKPG21-21 to Hua Peng, and support from Bioland Laboratory (Guangzhou Regenerative Medicine and Health Guangdong Laboratory) to Hua Peng.

## Author Contributions

Conceptualization, S.S., Y.C., H.P., and Y.-X.F.; Methodology, S.S, Y.C., T.S; Investigation, S.S, Y.C., T.S., Y.P., L.C., H.X., C.M., Y.L., J.S., S.Z., H.J., X.F., D.Y.; Software, S.S, Y.G., Y.C., Y.Z.; Resources, P.G., H.T., J.Z., Z.Z, J.Y., Z.H.; Writing-Original Draft, S.S., Y.C.; writing-Review-& Editing, S.S, Y.C., Y.P., H.P., and Y-X.F.; Supervision, H.P. and Y-X.F.; Funding Acquisition, H.P.

## Extended Figure Legend

**Extended Data Fig. 1:**
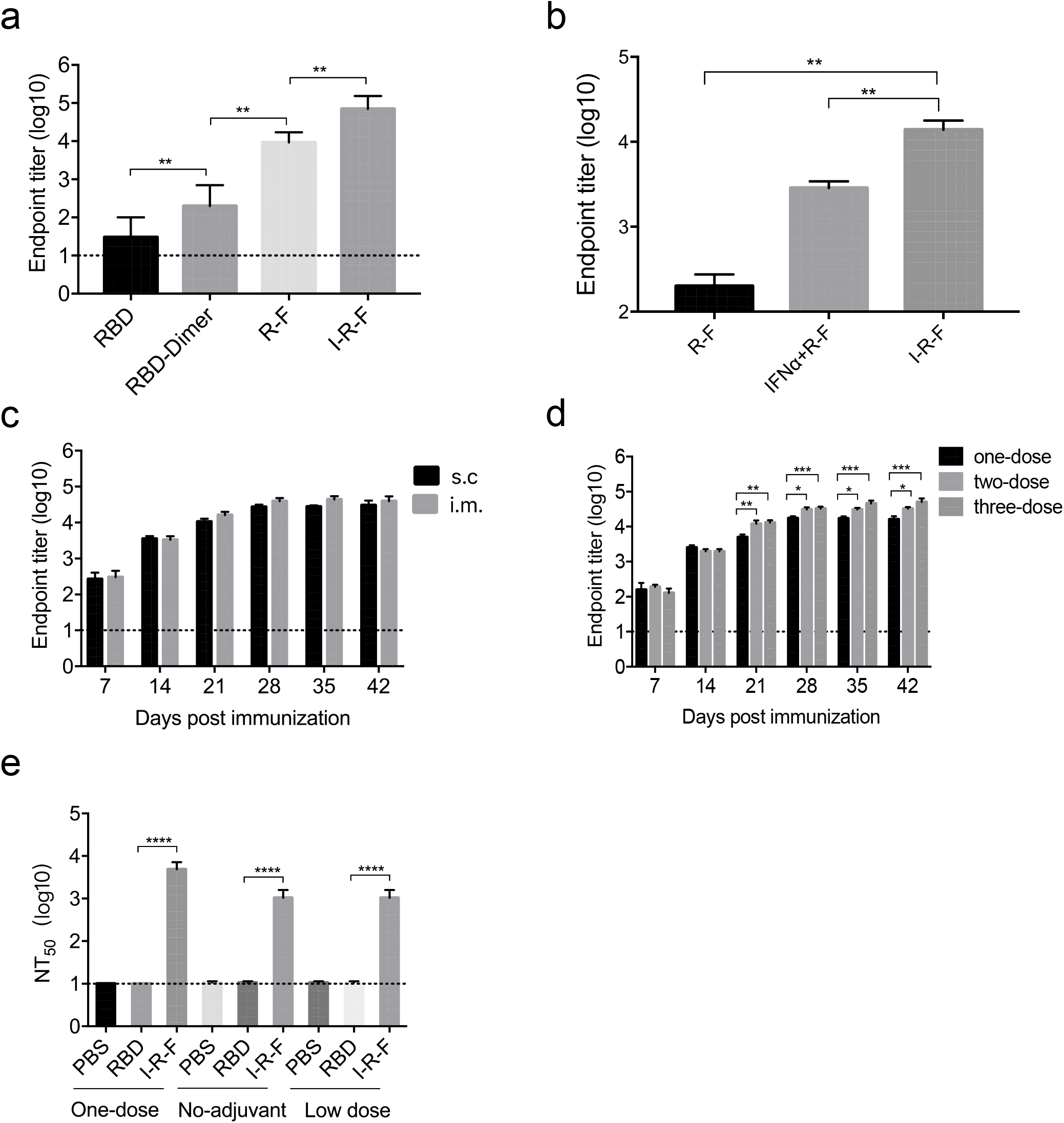
I-R-F induces more robust neutralization antibodies than RBD. (a). BALB/c mice (n=6/group) were i.m vaccinated with 10μg of I-P-F or equimolar of RBD, RBD-dimer, and R-F protein or PBS control with a signal vaccination regimen. The levels of RBD-specific IgG in serum on day 28 after immunization were determined by ELISA (b) BALB/c Mice (n=6/group) were intramuscularly vaccinated with 1μg of I-R-F or equimolar R-F plus IFNα protein without adjuvant and boosted with the same dose at a 14-day interval. Serum samples were collected on day 14 after the second immunization to evaluate the levels of RBD-specific IgG. (c) BALB/C mice (n=8/group) were immunized intramuscularly or subcutaneously with alum-adjuvanted 10μg of I-R-F and boosted on day 14 after initial immunization with equivalent dose. Serum samples were collected every week to determine the SARS-CoV-2-specific IgG antibody titers by ELISA. (d) BALB/C mice (n=8/group) were immunized intramuscularly with alum-adjuvanted 10μg of I-R-F by using a single dose (day0), two-dose (day0/14), and three-dose (day0/14/28) immunization procedures, respectively. Sera were collected on days 7, 14, 21, 28, 35, and 42 after the initial immunization and analyzed by ELISA to determine the IgG titer. (e) The neutralization antibody titers in serum described in Fig (2d-2f) on day 28 were determined by SARS-CoV-2 pseudovirus neutralization assay. The dashed line indicates the limit of detection. Data are shown as mean ± SEM. P-values were calculated by one-way ANOVA with multiple comparisons tests. ns (not significant), *P<0.05, **P<0.01, ***P<0.001, ****P<0.0001.

**Extended Data Fig. 2:**
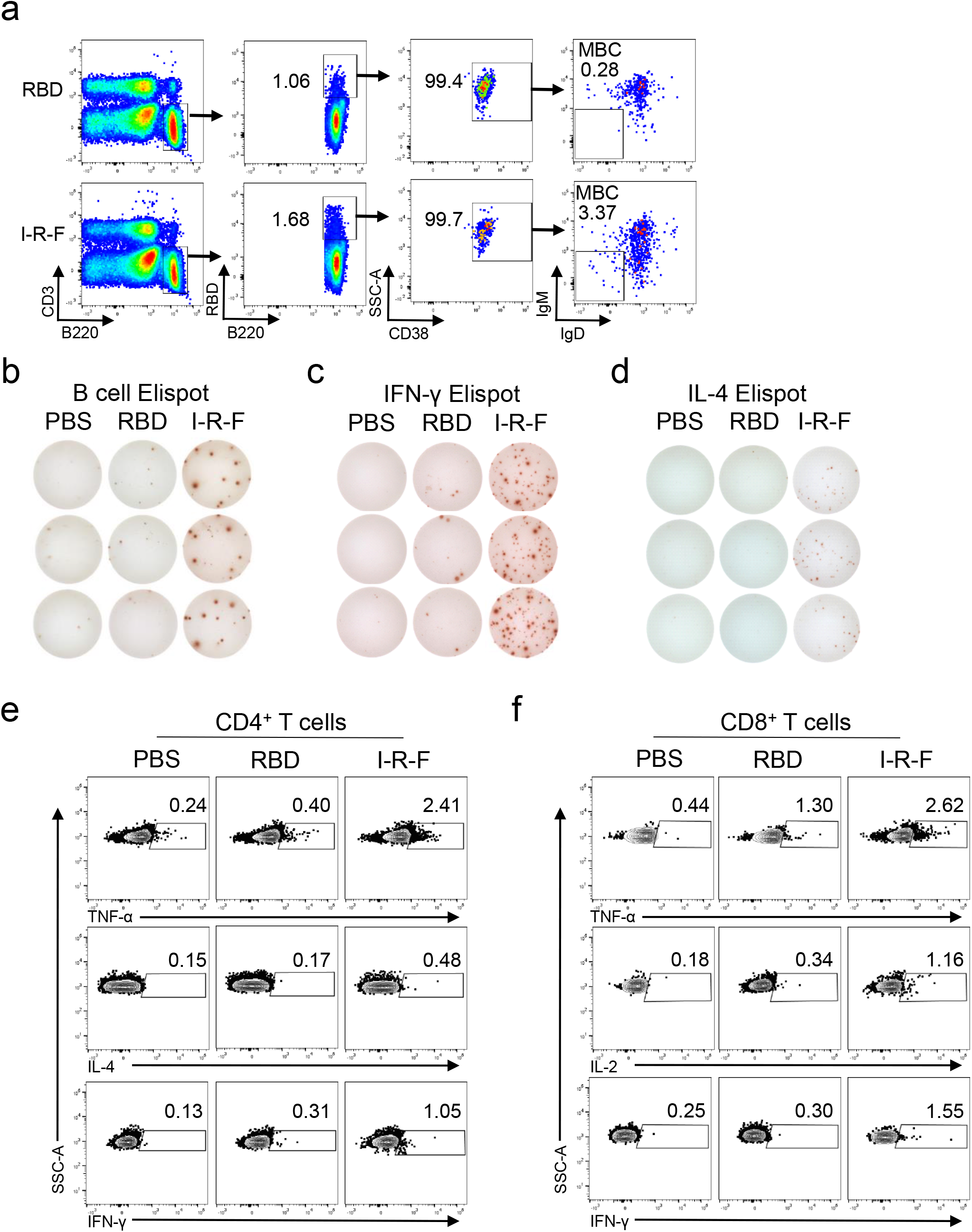
B and T cells immune response related to Figure 3. (a) Gating strategy for evaluating RBD-specific memory B cells in the spleen related to Fig.3c was present. (b-d) The raw data for B cell ELISPOT, IFN-γ, and IL-4 ELISPOT related to Fig3.b, d and e respectively were present. The representative flow cytometric gating related to Fig.3f, and Fig.3g was shown in (e) and (f).

**Extended Data Fig. 3:**
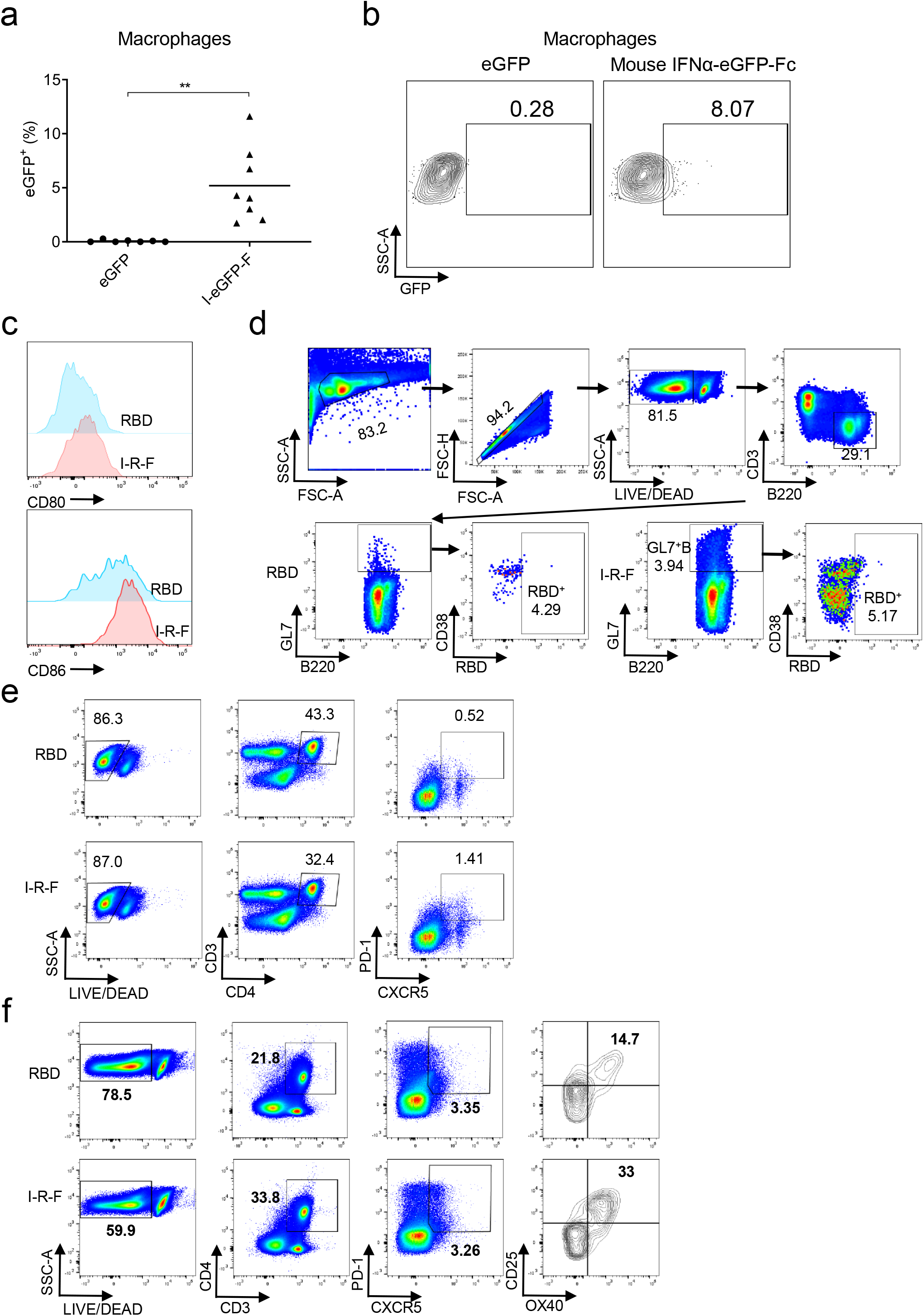
Potent antigen presentation, Tfh, and GC generation. (a and b) C57BL/6 mice (n=7 or 8/group) were intramuscularly injected 1nmole of I-E-F (mouse IFNα-eGFP-Fc) or eGFP. Four hours after injection, mice were sacrificed. Lymphocytes from mouse iLNs were collected to analyze the capture of I-E-F or eGFP. FCM analysis was proceeded to determine the percentages of GFP-positive macrophages (B220^-^CD11b^+^F4/80^+^). The results of GFP-positive macrophages (a) and the representative flow cytometric gating (b) were present. The representative flow cytometric gating related to Fig.4e is shown in (c). (d-f) Representative gating strategy for evaluating RBD-specific GC B cells, Tfh cells, and OX40^+^CD25^+^ Tfh cells related to Fig.4f-j.

**Extended Data Fig. 4:**
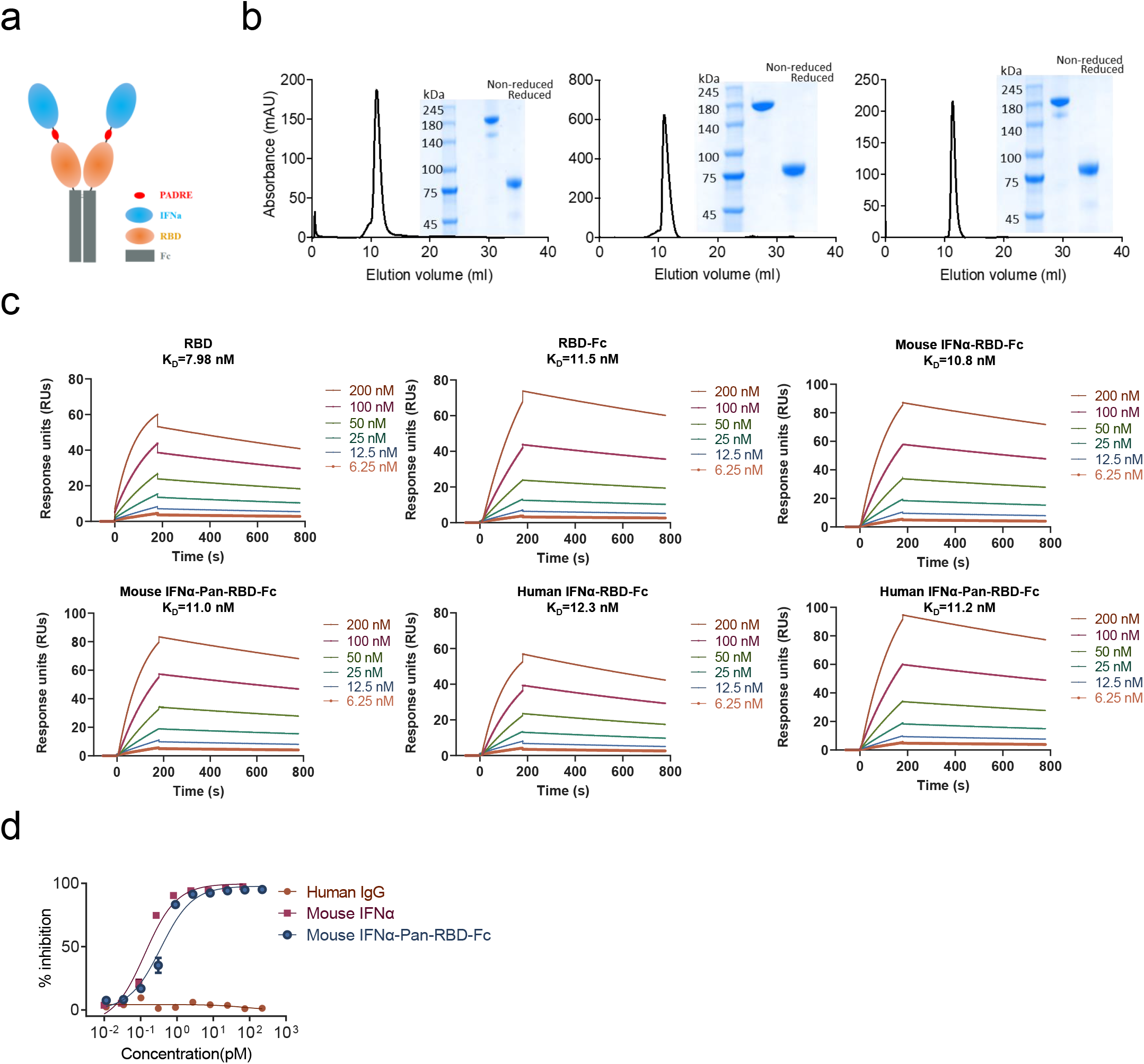
Design and characterization of the I-P-R-F (mouse), I-R-F (human), and I-P-R-F (human) vaccines. (a) The Schematic model of the I-P-R-F fusion protein. The showing structural elements contain a mouse IFNα4a (IFNα), (G4S3)_4_ linker (linker), Pan DR epitope (Pan), receptor-binding domain (RBD), immunoglobulin Fc fragment (Fc). (b) Size exclusion chromatography of mouse I-P-R-F, human I-R-F, and human I-P-R-F was performed on a Superdex200 Increase Column. The ultraviolet absorption at 280mm is shown. The insert pictures show the SDS-PAGE of the eluted samples. (c) Real-time association and dissociation profiles of RBD, RBD-Fc, mouse I-R-F, mouse I-P-R-F, human I-R-F, and human I-P-R-F bound to hACE2. The K_D_ value was calculated by the software BIA evaluation Version 4.1 (GE Healthcare). (d) The bioactivity of IFNα contained in mouse I-P-R-F was measured by an anti-viral infection biological assay.

**Extended Data Fig. 5:**
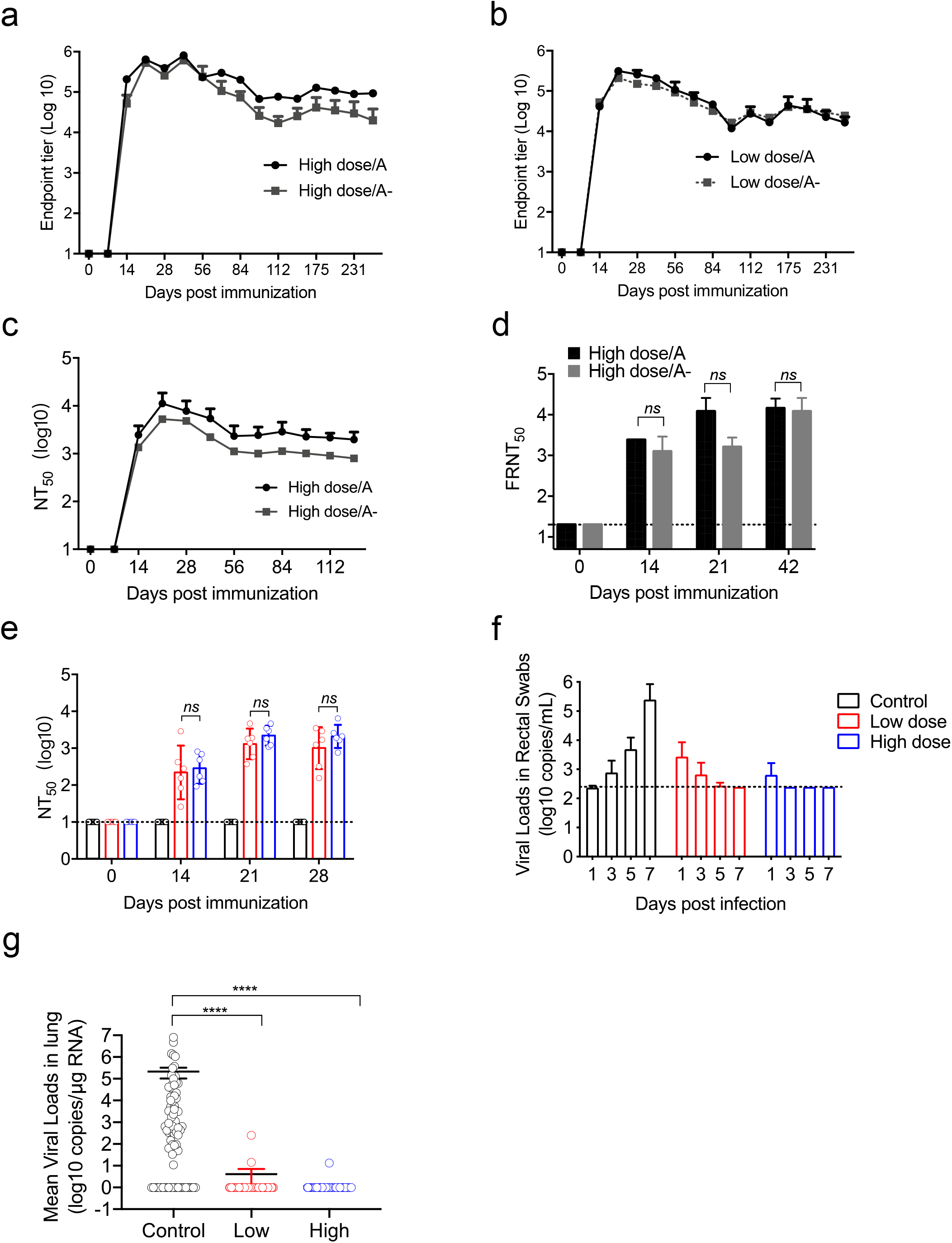
The immunogenicity and protective efficacy of I-P-R-F in rhesus macaques. Male and female rhesus macaques (n=8) were equally divided into four groups with a sex ratio of 1:1 and were immunized i.m. with a high dose (50ug) or a low dose (10ug) of I-P-R-F with or without alum as an adjuvant and boost with the same dose at a 14-day interval. The sera were collected every week are analyses for I-P-R-F immunogenicity and vaccine-induced neutralizing antibodies. **(a)** RBD-specific IgG titers in the high-dose group were determined by ELISA at indicated time points. **(b)** The dynamic IgG titers in sera from the low-dose vaccinated group were determined. **(c-d)** The kinetics of neutralization antibody titers in serum from I-P-R-F immunized animals were determined by SARS-CoV-2 pseudovirus and authentic virus neutralization assays. **(e)** Neutralization of SARA-CoV-2 pseudovirus by the anti-sera from I-P-R-F immunized rhesus macaques before the virus challenge. **(f)** Viral load in anal swabs of rhesus macaques challenged with live SARS-CoV-2. (g) Viral loads in the lungs. For each group of vaccinated macaques, 84 specimens from fourteen lung lobes were collected on day 7 post infection and subjected to determine the viral loads in the lungs. High dose/A: immunization with high does with adjuvant, High dose/A-: immunization with high does without adjuvant, Low dose/A: immunization with low does with adjuvant, Low dose/A-: immunization with low does without adjuvant. The dashed line indicates the limit of detection. Data are shown as mean ± SEM. P-values in (d) were analyzed with the unpaired t-test. P-values were calculated by one-way ANOVA with multiple comparisons tests in (e). ns (not significant), *P<0.05, **P<0.01, ***P<0.001, ****P<0.0001.

**Extended Data Fig. 6:**
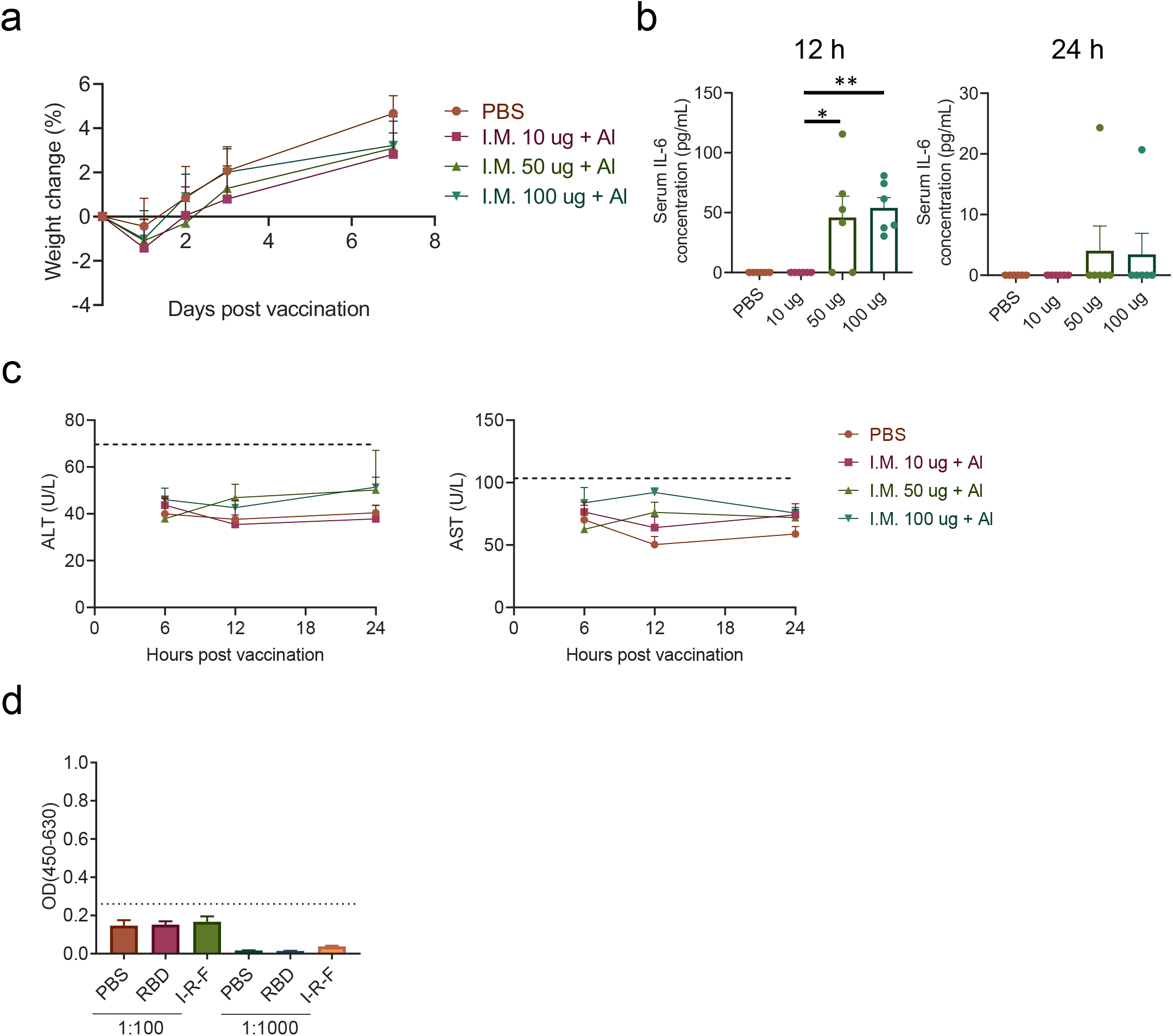
IFN-armed I-R-F induces no obvious toxicity. **(a-c)** Naïve female WT BALB/c mice (n = 6) were intramuscularly immunized with RBD or mouse IFNα-RBD-Fc, weighted, and bled. **(a)** Bodyweight changes were monitored and analyzed. **(b)** Serum inflammatory cytokines were determined by Cytometric Beads Array (CBA). **(c)** ALT and AST levels were analyzed by an automatic biochemical analyzer. The dotted lines indicate the upper limit of the normal range of ALT or AST using mean+2×SD of PBS group several hours after injection. **(d)** Evaluation of the IFNα specific antibody levels by ELISA. BALB/c Mice (n=6/group) were intramuscularly vaccinated with 10μg of I-R-F or equimolar RBD protein and boosted with the same dose at a 14-day interval. PBS was performed as a negative control. Sera were collected on day 28 after the initial immunization and subjected to determine the anti-murine IFNα level. The dashed line indicates the limit of detection. Data are shown as mean ± SEM. P-values were calculated by one-way ANOVA with multiple comparisons tests. ns (not significant), *P<0.05, **P<0.01, ***P<0.001, ****P<0.0001.

